# The antique genetic plight of the Mediterranean monk seal (*Monachus monachus*)

**DOI:** 10.1101/2021.12.23.473149

**Authors:** Jordi Salmona, Julia Dayon, Emilie Lecompte, Alexandros A. Karamanlidis, Alex Aguilar, Pablo Fernandez De Larrinoa, Rosa Pires, Giulia Mo, Aliki Panou, Sabrina Agnesi, Asunción Borrell, Erdem Danyer, Bayram Öztürk, Arda M. Tonay, Anastasios K. Anestis, Luis M. González, Panagiotis Dendrinos, Philippe Gaubert

## Abstract

Disentangling the impact of Late Quaternary climate change from human activities can have crucial implications on the conservation of endangered species. We investigated the population genetics and demography of the Mediterranean monk seal (*Monachus monachus*), one of the world’s most endangered marine mammals, through an unprecedented dataset encompassing historical (extinct) and extant populations from the eastern North Atlantic to the entire Mediterranean Basin. We show that Cabo Blanco (Western Sahara/Mauritania), Madeira, Western Mediterranean (historical range), and Eastern Mediterranean regions segregate into four populations. This structure is likely the consequence of recent drift, combined with long-term isolation by distance (*R*^2^ = 0.7), resulting from prevailing short-distance (< 500 km) and infrequent long-distance dispersal (< 1,500 km). All populations (Madeira especially), show high levels of inbreeding and low levels of genetic diversity, seemingly declining since historical time, but surprisingly not being impacted by the 1997 massive die-off in Cabo Blanco. Approximate Bayesian Computation analyses support scenarios combining local extinctions and a major effective population size decline in all populations during Antiquity. Our results suggest that the early densification of human populations around the Mediterranean Basin coupled with the development of seafaring techniques were the main drivers of the decline of Mediterranean monk seals.

## Background

Distinguishing between the respective impact of recent climate- and human-driven changes on the biosphere has proven challenging [2] because Late Quaternary extinctions were caused by the superimposed effects of climate change and anthropization from the Last Glacial Maximum (LGM) onwards (ca. 25–10 ka; [1]). Differentiating climate- and human-driven impacts is similarly relevant to currently endangered species. Indeed, informed conservation planning relies on a systemic approach including knowledge of species’ demographic history [2], which in turn can be used to predict species’ ability to adapt to future climate changes [3,4].

Pinnipeds are marine mammals that rely on coastal haul-out areas during their annual life cycle. As such, they have been affected by LGM climate changes and early human activities [5,6], although the impact of the latter (through targeted hunting) occurred after the end of the LGM in this case (but see [7]). Despite the fact that the Mediterranean monk seal (MMS; *Monachus monachus*) is arguably the world’s most endangered pinniped [8], the factors responsible for its critical conservation status are not well understood. The MMS once ranged across the entire Temperate Northern Atlantic province [9], from the Black Sea and the Mediterranean Basin into North Atlantic eastern waters encompassing the coasts of western Africa, the Macaronesian islands, and the northern Iberian Peninsula [10–12]. Nowadays, the species is fragmented into three isolated areas, distributed in the eastern Mediterranean (ca. 187-240 mature individuals [13]) and the eastern North Atlantic, in Cabo Blanco (Western Sahara/Mauritania; ca. 350 individuals) and the archipelago of Madeira (ca. 20 individuals; [14]) (Fig. 1).

**Figure 1.**
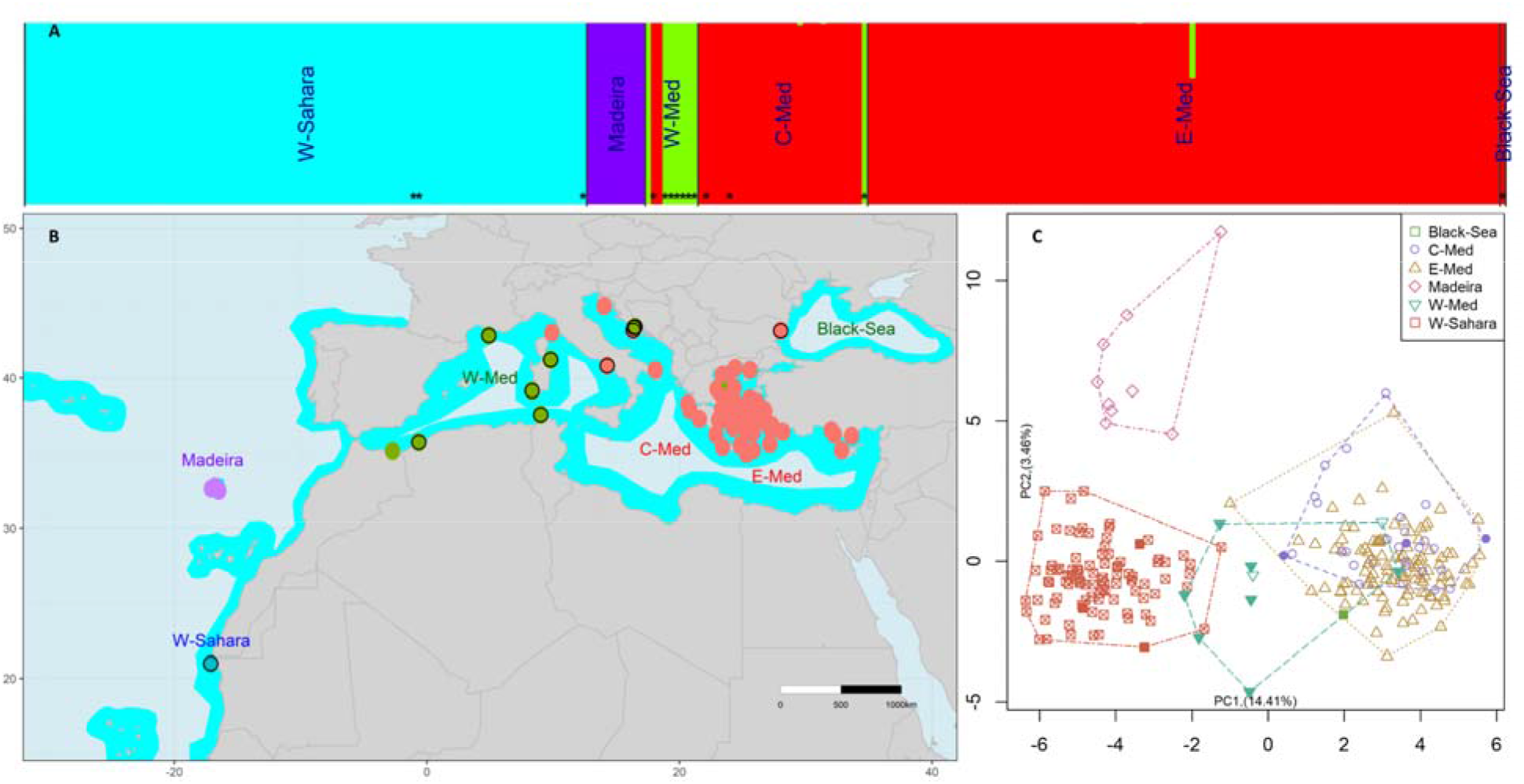
*Monachus monachus* nuclear genetic structure. Individual posterior cluster membership coefficients for *K* = 4 represented using vertical barplot (**A**) and geolocated pie-charts (**B**). Representation of the samples from the six main sampling areas on the first two axes of principal component analysis (**C**) showing a strong West-East cline of differentiation on the first axis. In **A**, the samples are ordered by regions, sorted from West to East, separated by vertical black lines, and stars denote historical samples. In **B**, black circles denote locations that include historical samples and the cyan area represents the putative ancient MMS distribution. In **C**, full and empty symbols represent historical and modern samples, respectively. W-Sahara: Cabo Blanco (Western Sahara/Mauritania), W-Med: western Mediterranean Sea, C-Med: central Mediterranean Sea, E-Med: eastern Mediterranean Sea.

A combination of extrinsic (human activities) and intrinsic (e.g., low genomic diversity, high susceptibility to diseases) factors have been proposed to explain the local extinctions affecting the species [15,16], and to a larger extent, the extinction proneness observed in the entire Monachinae subfamily (which also includes the endangered Hawaiian monk seal *Neomonachus schauinslandii* and the extinct Caribbean monk seal *N. tropicalis* [17]). As an iconic representative of the Mediterranean Basin biodiversity hotspot [18], the MMS could have suffered from targeted hunting since the advent of the early Mediterranean sailors (Bronze Age, ∼3,300 to 1,200 BCE; [19–21]). From the end of the Middle Age (15^th^ century), the massive exploitation of the species—notably in the Atlantic Ocean—became documented through the logbooks of European marine explorers [11,22]. Over the last centuries, MMS populations have been further impacted by the expansion of the fishing industry, deliberate killing by fishermen, marine pollution, and human coastal encroachment, leading to local extinctions in most of the species’ range, notably in the western Mediterranean and the Black Sea [12,23–25].

The decline of the MMS is documented from historical mass killing [10,19,26], local extinctions [27], massive die-off [28], and genetic inferences [29]. However, the respective impacts of Late Quaternary climatic fluctuations and anthropogenic pressures on the demographic history of the species remain poorly understood. Yet, the LGM induced significant changes in the Mediterranean Basin, such as lowering of sea level (down to -120 m; [30]) and sea surface temperatures, and local variations in salinity [31,32]. Moreover, the impacts of human activities on MMS populations may have applied at different periods across the species’ range, with early exploitation since the Middle Paleolithic [20,33], followed by a gradual dispersal of seafaring civilizations from the eastern Mediterranean Basin (where coastal encroachment and marine resources’ exploitation have been documented since ca. 10,000 yrs) towards the western Mediterranean Basin and North Atlantic Ocean [21,30,33,34].

Previous genetic studies have revealed the genetic isolation of the three extant populations of MMS and their low levels of genetic diversity (since at least the mid-19^th^ century), and the local extinction of mitochondrial DNA (mtDNA) haplotypes in the western Mediterranean Basin [35–38]. Genetic data additionally disclosed a demographic bottleneck in the Cabo Blanco [29,39], and a signature of past metapopulation dynamics across the species’ range [37,40]. In this study, we investigate the genetic patterns and demographic history of the MMS through an unprecedented dataset, including historical individuals collected across the majority of the species’ range and the three extant populations. We use microsatellite genotyping and mtDNA sequencing of 375 MMS to assess the historical population structure and dynamics of the MMS across the Mediterranean Basin and Northern Atlantic Ocean. Specifically, we delineate past and present population structure and diversity and assess whether such patterns are consistent with scenarios of local extinctions together with genetic drift and inbreeding in extant populations. We also investigate the different historical drivers (LGM *vs*. human activities) potentially affecting the MMS population dynamics across the Mediterranean Basin and Northern Atlantic Ocean through complementary population genetic and modeling approaches based on demographic models, incorporating population structure and connectivity among populations through time. Based on our results, we formulate recommendations that might contribute to future evidence-based conservation strategies of the MMS.

## Methods

### Laboratory procedures and genotyping

We collected 383 samples from recent (N=314; 1989 – 2020) and historical (N=69; 1833 – 1975) specimens of MMS covering the extant populations’ and historical species’ range (Figs 1, S1-3), within the frame of the authorizations, and of the best practice guidelines listed in Method S1. The delineation between recent (from 1989 on) *vs*. historical (prior to 1975) is based on the extinction period of the last resident MMS groups from the western and parts of the central Mediterranean Sea during the 1970s [19], which suggested that after this period the species distribution fragmented into North Atlantic and eastern Mediterranean populations.

Genomic DNA was extracted from fresh tissue and skin samples, hairs, feces, bones, and tanned skin using dedicated protocols (Method S1). We amplified 524 bp of the hypervariable region I of the mitochondrial control region (CR1) following [41] for the modern samples (N = 121), and reconstructed 484 bp fragment encompassing all the variable sites of CR1 for the historical samples (N = 3), as already described elsewhere [35] and detailed in method S2. We complemented our original dataset with 209 already published CR1 sequences [35,36,41]. Final CR1 alignment included 232 modern, 23 dated historical samples, and four historical samples with unknown dating, for a total of 326 sequences (Figs 1, S1-3, Tables S1-3).

We genotyped 383 samples at 19 nuclear microsatellite loci ([38], Method S3) from 314 modern samples and 69 historical samples. In order to mitigate scoring errors and potential allelic dropout, PCRs of DNA extracts from hairs, feces, and museum material were systematically replicated 2-to-5 times ([38], Method S3). We controlled markers for allelic dropout, linkage disequilibrium and departure from Hardy – Weinberg equilibrium (Method S3, Table S4; Fig. S4-7), and applied a two-fold sample selection procedure to optimize the number of retained individuals genotyped at informative loci. We relied on the minimum number of loci necessary to discriminate among individuals and the region-based discriminant analysis of principal components [DAPC] contribution of loci (Method S3, Fig. S8). The final microsatellite data set comprised 253 samples (including 14 historical and 239 modern samples) with 10.18 % missing data (Tables S1-2, Figs 1 & S1-2).

### Genetic diversity and structure

To assess the overall, per sampling site, and per locus microsatellites genetic diversity, we estimated the allelic richness (*A*_R_ [42]) with the R package *hierfstat* [43]. We also assessed the number of alleles (*A*), the observed (*H*_O_) and Nei’s unbiased expected heterozygosities (*H*_E_ [44]) with the R package *adegenet* [45]. To evaluate the effect of drift in the small colonies of MMS, we estimated the average individual inbreeding coefficient *F* [45] in *adegenet*, from 100 iterations. We assessed the level of genetic differentiation among localities, populations, and time periods using Nei’s *F*_ST_ [46]. Additionally, we investigated patterns of genetic variance with a principal component analysis (PCA) of allele frequencies, with a discriminant analysis of principal components (DAPC, considering sampling regions as groups), and using the SnapClust clustering approach [47] for *K* = 1-15. To assess how the geographic distance alone explains the genetic diversity [48,49], we investigated individual- and population-based patterns of isolation by distance (IBD) using Mantel tests [50], and assessed which genetic distance metric and spatial scale best fitted the data [51,52]. To assess how dispersal is distributed geographically we used Mantel correlograms [53,54].

For CR1, we estimated the number of haplotypes (*Nh*) and polymorphic sites (S), haplotype diversity (*h* [55]), and nucleotide diversity (π [56]) per sampling area using the R package *pegas* [57]. CR1 haplotype relationships were reconstructed with a maximum parsimony network using *pegas*. All the above-mentioned methods are detailed in Method S4.

### Demographic history

We used Approximate Bayesian Computation (ABC; [58]) to compare data simulated under several alternative scenarios to the real data (microsatellites & CR1), and estimate parameters of interest from the best-supported scenario [59–61]. We first tested scenarios assuming a single panmictic population and different histories of population size change (Figs S9-10). Second, we modeled stable-size structured populations with constant migration rates among populations (Figs S9, S11). These models included unsampled ghost populations mimicking extinct populations and a change in connectivity, mimicking the loss of gene flow among extant populations (Figs S9, S11-12). Third, we combined population size change, structure, ghost populations, and changes in connectivity, to model populations that suffered one or several events of decline and fragmentation (Figs S9, S13-14). Structured scenarios also compared n-island, stepping-stone, and spatially-explicit models of connectivity (Table S5, S6). To overcome the potential overfitting of parameters, we reduced the summary statistics dimension [62], both using partial least square regression (PLS) and minimizing the sample entropy [63]. We retained models with low marginal density (MD), high proportions (MD *p-value*) of retained simulations showing a lower or equal likelihood under the inferred GLM as compared to the observed genetic data [59], and with high centrality of the observed data within the multidimensional cloud of retained simulations (Tukey *p-value* [64]). We assessed the ABC’s ability to distinguish between the proposed models, with 1000 pseudo observed data sets (pods) randomly selected from simulated data sets under each model. All the above-mentioned methods are detailed in Method S6.

## Results

### Genetic diversity and structure

The microsatellite population-structure analyses exhibited a strong East-West pattern of differentiation (*F*_ST_ > 0.4, Table S7-8) separating the Atlantic and Mediterranean populations into four clusters (Figs 1, S15-16). This pattern appeared as a continuous cline of differentiation driving the first component of the PCA (Fig. 1) and was also the first revealed by the DAPC (Fig. S17). Second, our analyses showed clear segregation of Madeira from Cabo Blanco individuals, distinguishing the two Atlantic populations (Figs 1, S16-17). The Western-Mediterranean population formed a group relatively distinct from the Eastern-Mediterranean and was genetically intermediary between the former and Atlantic populations (Figs 1, S16-17). The only historical sample genotyped from the Black Sea was assigned to the Eastern-Mediterranean population (Figs 1, S16). Further subdivision of the MMS resulted in erratic clustering results (Figs S15-16, S18).

Overall, all the populations showed relatively low nuclear genetic diversity (*H*_O_ = 0.12 - 0.42 & *H*_E_ = 0.04 - 0.36) and a wide range of inbreeding levels (Fig. 2). Madeira had the lowest levels of genetic diversity across all estimated indexes and among the highest estimates of inbreeding (Fig. 2, Tables S7-8). The historical Western-Mediterranean population showed a relatively high allelic richness, and the Cabo Blanco and Eastern-Mediterranean populations exhibited the highest number of private alleles (Fig. 2). The historical sample from the Black Sea did not harbor private alleles (Fig 2). Similarly, mtDNA (CR1) diversity was low in all populations, with Madeira exhibiting only one haplotype (Table S3, Fig S19).

**Figure 2.**
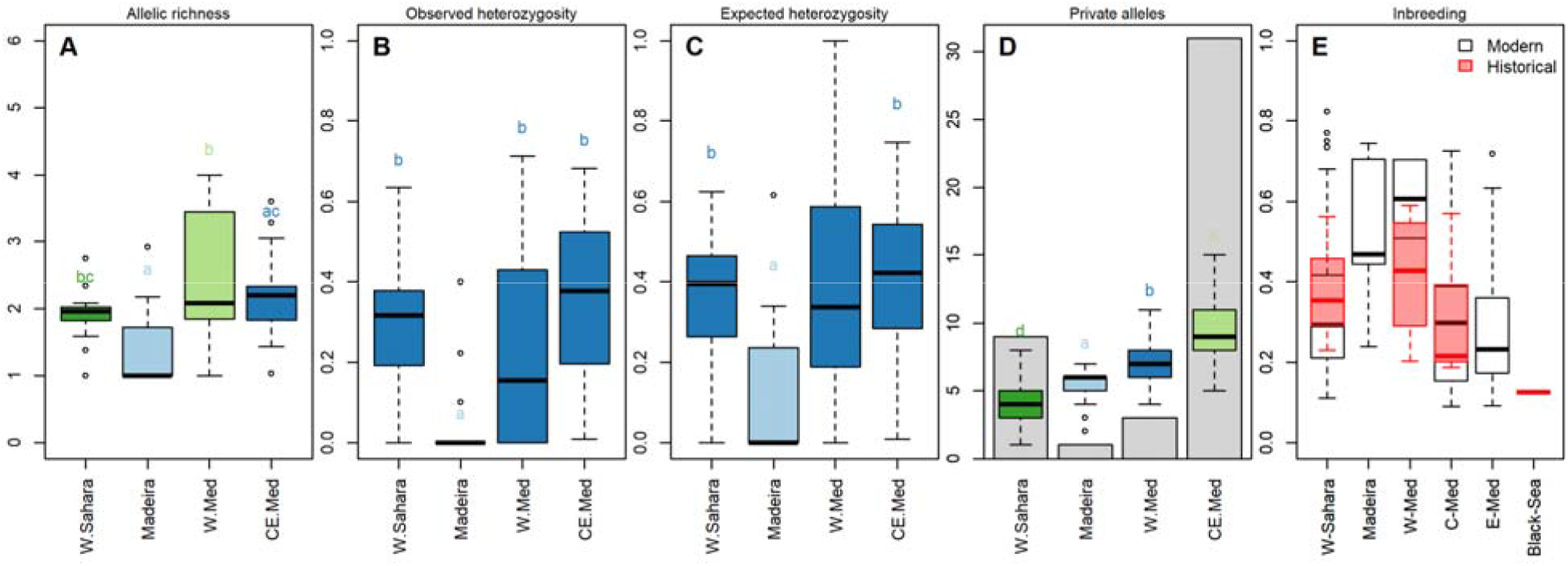
*Monachus monachus* nuclear genetic diversity. Boxplot of the microsatellites (**A**) allelic richness (*A*_R_), (**B**) observed (*H*_O_), and (**C**) expected (*H*_E_) heterozygosity, (**D**) mean private alleles (*P*_A_) of 500 resampling of the smallest sample size across each sampling area, and barplot of the number of private alleles (*P*_A_), and (**E**) boxplot of the average individual inbreeding coefficient (*F*), per sampling area, expressed for modern (>1979) and historical (<1980) samples. In A-D, historic samples were removed from the analyses for all extant populations apart from the W-Med where no population is clearly identified and for which we almost exclusively have historic samples. Letters and box colors in A-D illustrate the Tukey posthoc group assignment. W-Sahara: Cabo Blanco, W-Med: Western Mediterranean Sea, CE-Med: merged Central and Eastern Mediterranean Sea, C-Med: Central Mediterranean Sea, E-Med: Eastern Mediterranean Sea.

The likelihood estimates of individual homozygosity (*F*) suggested higher levels of inbreeding in modern samples than in historical ones (per population; Fig. 2). In addition, the overall estimates [all historical *vs* all modern samples] of allelic richness (*A*_R_) and private allelic richness (*P*_A_) were higher historically than in modern samples (Fig. S20). However, the overall observed (*H*_O_), and expected (*H*_E_) heterozygosity, as well as the overall evolution of inbreeding levels through time (Fig. S20), did not exhibit a pattern of diversity loss across the timespan covered by our dataset (1840-2020). Across modern samples (>1975), and for the two major populations (Cabo Blanco and Eastern-Mediterranean), we could not see any clear pattern of diversity loss between 1990 and 2020 (Figs S21-22). At the mtDNA level, one historical CR1 haplotype was not recovered in modern samples (MM07) and, reciprocally, one modern haplotype was not recovered from historical samples (MM06; Figs S19, S23-24).

In 1997 the Cabo Blanco colony underwent a massive die-off (over two thirds of the population was wiped out), caused either by saxitoxins or by a morbillivirus outbreak [28,65,66]. Our results based on various population and individual-based genetic diversity indexes (*A*_R_, *H*_O_, *H*_E_, *F*), apart from private alleles (*P*_A_), were not consistent with the genetic diversity decrease expected after a bottleneck despite a large sampling before and after the event took place (except for private alleles [PA]; Figs S25-26). Furthermore, we did not record signals of rare allele loss, or decrease in frequency, nor of major allele gain in frequency (Fig. S27-28), expected after a reduction of the population size [67,68].

In line with the PCA showing an East-West cline of differentiation, we found an isolation by distance (IBD) pattern explaining up to 69.6% of the among-individuals genetic distance variability (Figs 3, S29-30). Such a strong IBD signal was sustained by Mantel correlograms exhibiting high values for the first 500 km classes, progressively declining up to ∼1,500 km, above which values were no longer significant or negative (Fig. 3). Females exhibited significant positive values up to higher distance classes (1500 km) than males (700 km), but this signal may have resulted from sample size differences among males and females for each class.

**Figure 3.**
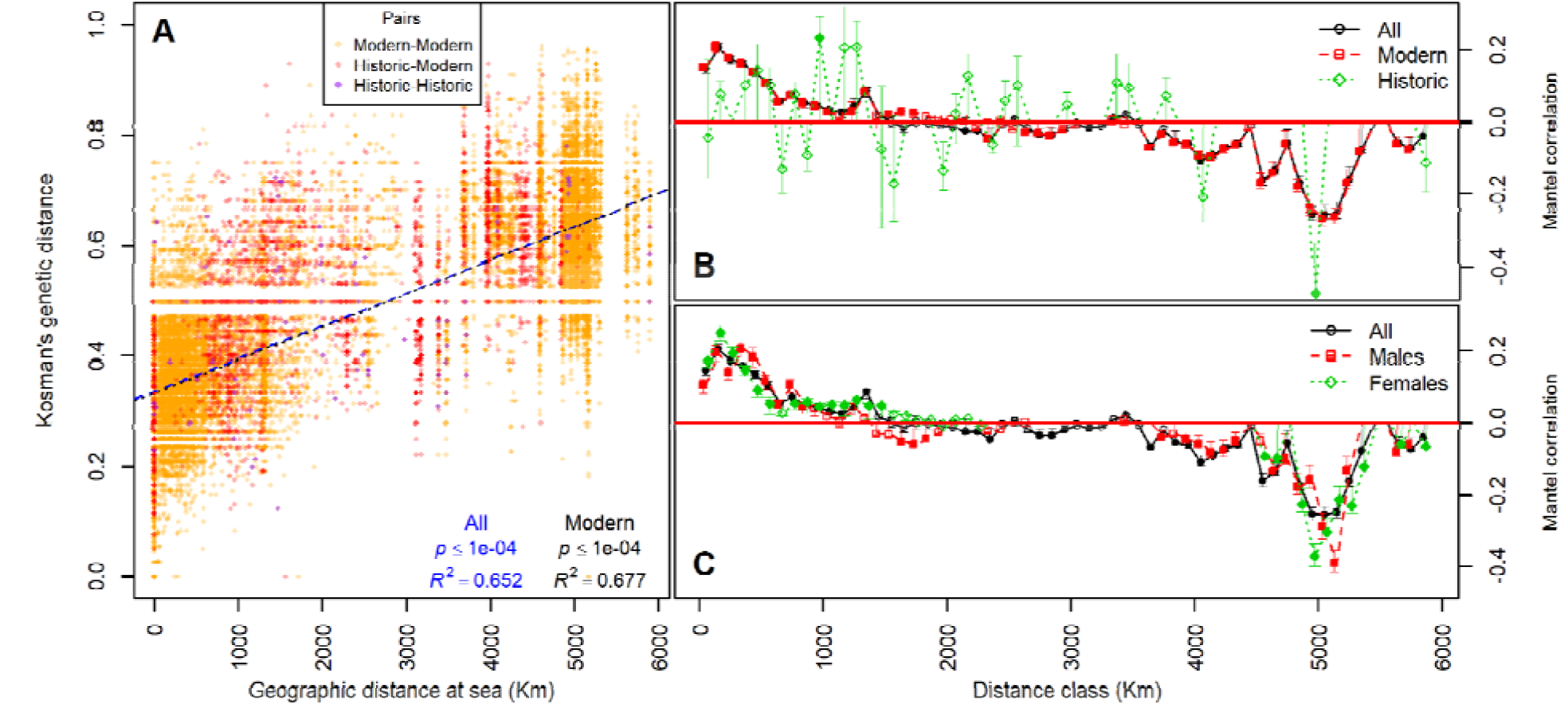
Isolation by distance in *Monachus monachus*. **A**: isolation by distance (IBD) relationship between individual-based genetic and maritime distances. **B** & **C**: Mantel correlogram of spatial correlation, for 59 classes of 100 Km, in all, modern, and historical samples (**B**), and all, males and females samples (**C**). Significant values, over 1000 permutations, are represented by filled symbols. Complementary analyses using alternative genetic distances are presented in Figs S27-28.

### Demographic history

The ABC procedure supported models including population structure, ghost populations (local extinctions), and an effective population size decline (Figs. S9-13). The most supported population structure used a custom stepping stone framework, realistically modeling extant and extinct population connectivity, based on their respective position in space (Table S4, S9-10, & Figs S12-14). Among the models including these features, the most supported ones (M180 & M181, Fig. S14) suggest that all sampled populations underwent one to two major declines, most of which were of at least one order of magnitude (Table S11, Figs 4, S31). Furthermore, the best models reveal that the declines, and the loss of connectivity among sampled populations, most likely occurred during Antiquity, and at the onset of the Middle-Age (Figs 4, S31-33).

**Figure 4.**
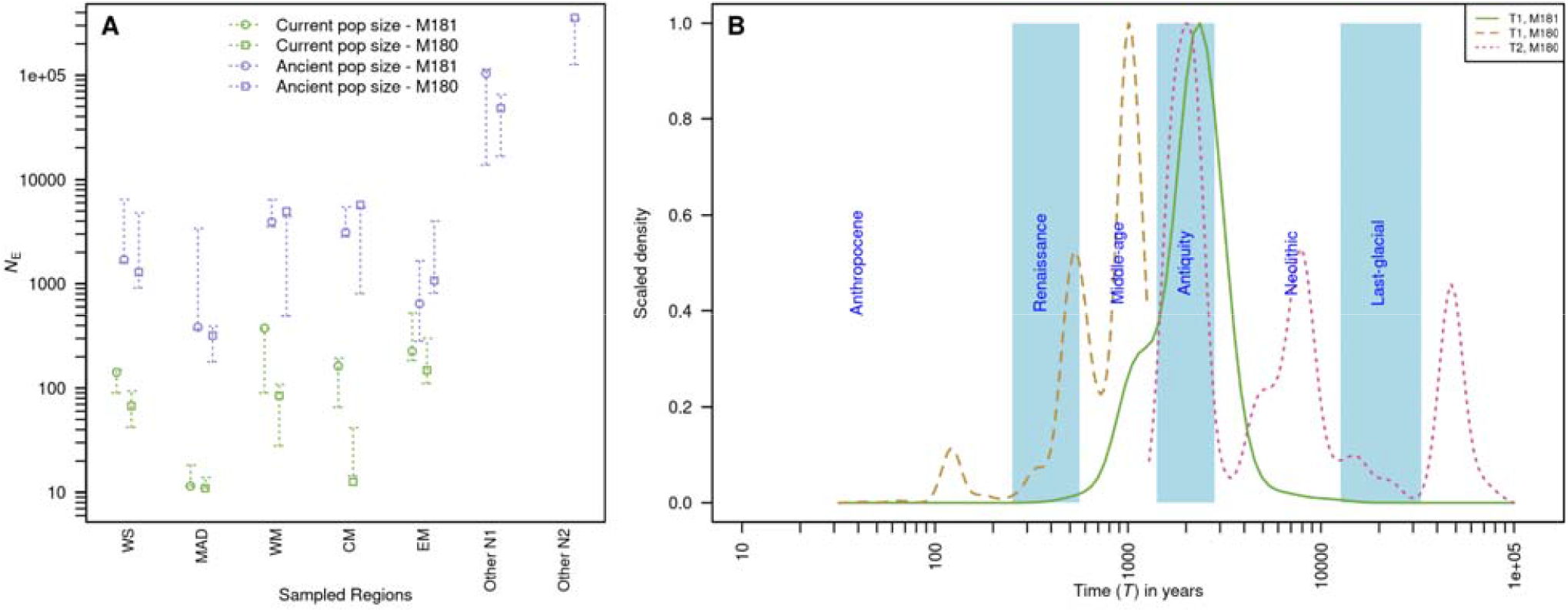
The antique decline of MMS. ABC-GLM posterior estimates of effective population size (*N*_e_; **A**) and of the time (*T*_1_ and *T*_2_; **B**) of the major demographic event(s) - Population decline + local extinction + change in connectivity - estimated from the most likely scenarios (M181 & M180), within the ABC framework. **A**: mode (symbol) and 50 % quantiles (dotted arrows) of the effective population size (*N*_e_) under the models 181 and 180, represented after (*N*_0_: current population) and before (*N*_1_ and *N*_2_: ancient populations) the demographic events occurring at *T*_1_ and at *T*_2_. **B**: scaled-density distributions of the time of the demographic event(s) *T*_1_ and at *T*_2_. WS: Cabo Blanco, MAD: Madeira, WM: Western-Mediterranean, CM: Central Mediterranean, EM: Eastern-Mediterranean, Other N1 and Other N2: cumulated ancient effective population sizes of other populations before *T*_1_ and *T*_2_ respectively.

## Discussion

### Genetic structure of Mediterranean monk seal through time and space

Our study is based on an unprecedented genetic sampling, covering the entire distribution range of the Mediterranean monk seal and encompassing historical (extinct) and extant populations. This enabled us to provide the most comprehensive diversity and structure assessment of the species to date, in comparison to previous studies limited in their geographic and temporal representation and/or genetic information. Our results illustrate how the inclusion of historical samples helps to understand the dynamics of species’ genetic diversity through time and space [69], despites potential allelic dropout associated with microsatellite genotyping of historical samples [70]. The global analysis showed that MMS are composed of four populations, including the previously delineated eastern Mediterranean (reaching the Ligurian Sea at its westernmost location) and two northern Atlantic populations (Madeira and Cabo Blanco) [35–38,41], and a newly identified, historical population in the Western Mediterranean. The latter ranged from the northern Maghreb to southern France, Sardinia Isl. and the Adriatic Sea, partly overlapping at its eastern fringe with the extant eastern Mediterranean population (Fig. 1). Because the western Mediterranean population holds both North Atlantic and Mediterranean mitochondrial haplotypes, it could not be clearly delineated in the previous, single-locus studies integrating historical samples [35,36]. The modern sample from northern Morocco (1993) raises the long-standing question as to whether the western Mediterranean population still exists today, notably on the western Algerian coast where MMS were present until the recent past [71]. Although sightings of MMS across the western Mediterranean basin have been reported within the last c. 20 yrs [72], their actual relevance is difficult to assess unless proper monitoring of potential breeding sites is systematized.

The status of the extinct population in the Black Sea [73] remains unsolved. The sole successfully genotyped historical sample included in the microsatellites analyses did however carry information attributing it to the eastern Mediterranean population (Figs 1, S16). Together with the sharing of a unique CR1 haplotype in the Black Sea with individuals from the Aegean Sea, our and previous results [35,36] do not support the straits of Çanakkale (Dardanelles) and Istanbul (Bosporus) as a putative barrier to MMS movements, despite its effect on other species at a wide taxonomic scale, including marine mammals [74,75].

Although substantial differentiation was previously reported between Cabo Blanco and Madeira [38], Cabo Blanco and Eastern Mediterranean [37], and Aegean and Ionian seas [40], our results demonstrate that this differentiation is clinal and likely a consequence of isolation by distance (*R*^2^ = 0.7, Figs 1, 3). At the level of the Mediterranean basin, similar continuous differentiation is found in several marine organisms with varying mobility [e.g. dolphins, Sea star [76,77]], suggesting that oceanic distance alone can mold species genetic diversity across a broad taxonomic range. Because the inclusion of historical samples did not disrupt the genetic covariation with geography, the isolation by distance pattern might have been long-standing over the past two centuries. Interestingly, this pattern, previously unveiled at the eastern Mediterranean Sea scale [40], appears driven by prevailing short distance (< 500 km) and infrequent long distance dispersal (< 1,500 km, Fig. 3). MMS may tend to establish in proximity to their place of birth, eventually dispersing within their local water basin, and, on rare occasions, to further-away places. This is in sharp contrast with the lack of structure of *Neomonachus schauinslandi*, the Hawaiian Monk Seal [78], which shows long distance dispersal across a ∼2,700 km linear oceanic distribution. On the contrary, our results suggest that natal philopatry, exacerbated by the rarity of suitable habitat and breeding sites, and by the weaker and circular Mediterranean Sea currents, are likely contributors to the current MMS genetic structure [79,80].

### A long depauperate and decreasing genetic diversity

Combining data from all extant populations, our genetic survey confirms that MMS harbors the most depauperate genetic diversity (*H*_E_ = 0.04-0.36) of all seal species [29]. Furthermore, it reveals Madeira’s dramatic levels of genetic diversity (*H*_E_ = 0.04, Fig. 2), which echoes its decline to less than 10 individuals in the 1980s [26]. As a matter of comparison, Madeira’s diversity is lower than that of the ringed seal population (*Pusa hispida*) landlocked since the LGM in Saimaa Lake, Finland [81].

We uncovered a decrease of allelic richness (*A*_R_) and private allelic richness (*P*_A_) from historical to modern samples, and an increasing trend in inbreeding at the population level (Figs 2, S20). This pattern is consistent with the trend exhibited by CR1 [35] and mitogenome [36] studies, confirming that MMS genetic diversity has decreased over the past two centuries. In all cases, the decrease in genetic diversity started from remarkably low historical levels, which could explain the relatively stable distribution of heterozygosity that we observed over time (Fig. S20). This contrasts with the sharp drop in diversity caused by commercial exploitation in other seal species [82,83], implying an older exploitation of MMS populations.

Remarkably, the MMS genetic diversity has not been decreasing in the past 30 years (Figs S21-22), at least in the two populations with sufficient temporal sampling (eastern Mediterranean and Cabo Blanco). This encouraging signal can be interpreted as the incipient results of conservation efforts and population recovery over that period [12]. However, this pattern is surprising in the Cabo Blanco, where we could not trace any substantial genetic diversity or minor allele frequency drop - except private allele loss - following the massive 1997 die-off that decimated more than two thirds of the Cabo-Blanco colony (Figs S25-26 [28]). The originally low diversity and high inbreeding in the population, a higher death rate of most inbred individuals at the time of the die-off, or an insufficient resolution of the studied loci, may have blurred declining pattern signatures. Additionally, individuals from unsurveyed nearby localities (e.g. Guerguerat) at the time of the mass die-off may have participated in maintaining the genetic diversity of the Cabo-Blanco colony (pers. obs. AA).

### Human-seal interactions in the Mediterranean basin: a history of overexploitation

The low diversity and high levels of inbreeding in all MMS populations (Fig. 2) are strong signals for the species undergoing a major decline with limited gene flow among populations. Furthermore, the marked IBD pattern (Fig. 3) reveals that MMS populations were connected by gene flow in the recent past. These patterns are confirmed by our demographic modeling analyses, which suggest that populations were organized in a stepping-stone manner, connected by intermediary (extinct) populations. These populations underwent one to two major demographic declines during Antiquity [∼800BC-600AD], most of which were at least of one order of magnitude (Figs 4, S31).

Surprisingly, the LGM (∼20 ka ago) that dramatically affected the sea level (∼120-130 m below current level) and the Mediterranean basin area, did not leave an identifiable signature in the MMS genetic data. One of our two best models also shows weak support for a population size decrease during the Neolithic [∼12,000 - 800 BC; Fig. 4]. The Neolithic transition allowed rapid human population growth and the development of complex civilizations [30,84], mastering increasingly sophisticated seafaring and fishing techniques [85,86]. By the rise of Antiquity, large human populations had spread across the entire Mediterranean basin and its islands [86–88]. Although the extent of hunting pressure on MMS is hard to accurately gauge at this time, historical sources relate hunting, meat consumption, oil use in lamps and skin use [89], conflict with fishermen [90], use in circus shows [91] and use of body parts to produce medicines [89]. In Antiquity, the species was reported to be common, widespread, and of ‘naive’ behavior towards humans, with rookeries of large size, and using open environments such as beaches, outcrops, or promontories [89,92,93]. Such descriptions echo the large ancient effective population sizes (between 100,000 [*N*_1_-M181] and 356,000 [*N*_2_-M180] individuals, Table S11) inferred from our demographic reconstructions, and are consistent with the ancestral population size estimates of *Neomonachus tropicalis* [94]. However, it is in sharp contrast with the current elusive nature of the species, its habits of resting and giving birth in caves and remote islets, all of which are probably a result of long-lasting persecution [8].

Whether Romans, as previously posited [10], and/or other ancient civilizations played a significant role in the abrupt demographic decline of MMS remains unsolved, given the lack of direct evidence. However, the Roman Empire’s large-scale wildlife exploitation that led to local-fish stock depletion and local megafauna extinction [91,95,96], and the emerging evidence of their whaling activities [97] are clues pointing towards their potential role in the demise of MMS. Importantly, humans have been hunting MMS long before Antiquity [20,33,98,99] and have continued afterward [11], together with increasing competition for habitat use and marine resources. Therefore, the MMS decline was likely a continuous process, with a peak during Antiquity that dramatically sealed the genetic impoverishment of the species. Furthermore, sealed animals were undoubtedly processed on the beach, and this explains that only very few bones reached inland sites [99]. Thus, systematic sealing (like whaling [97]) is unlikely to have produced large archeological accumulations, as is the case for smaller marine resources, like tuna [100]. The resulting paucity of ancient MMS remains [98,99] limits our ability to accurately conclude a precise period of MMS overexploitation.

### Conservation outcomes

The existence of a western Mediterranean MMS population, likely on the verge of extinction, calls for the urgent identification and protection of breeding areas. Prior sightings suggest that several sites potentially hosting individuals belonging to this population should be surveyed, monitored and adequately protected on priority (e.g. from Al Hoceima to Cap des Trois Fourches - Morocco, Balearic Islands - Spain, Corsica -France, Sardinia, Tuscan Archipelago, Sicily - Italy, La Galite - Tunisia [72,101]).

The clinal pattern of genetic diversity across the entire species distribution supports the absence of locally manageable conservation units. It, therefore, calls for an ambitious integrative conservation plan that comprehensively includes all countries (30) of the MMS past distribution range. Such concerted actions should be strengthened by governmental and intergovernmental agencies’ efforts and policies [102].

The low genetic diversity of extant populations calls for the development of an ambitious program that includes (i) increasing the size of colonies and populations and (ii) restoring genetic diversity and connectivity among extant populations. Ensuring population growth should be at the core of the MMS conservation program, as it will directly improve the resilience of populations to stochastic events [28]. Ways of re-establishing dispersion among populations should be considered, through, for example, the conservation of historical breeding sites (see [72]).

However, the dramatic levels of genetic diversity reported from our study indicate that rescuing the genetic diversity of MMS through the translocation of individuals among extant populations [103,104] is a scientifically-backed option to promote the long-term recovery of the species and should not be disregarded. Translocation should balance the benefits of counteracting the present limited gene flow among populations with any potential detrimental effects, such as the introduction into the receiving MMS community of pathogens to which it had not been previously exposed.

Furthermore, our results demonstrate that all extant MMS populations were connected until recently, and that their differentiation is explained by their frequent short distance (< 500 km) and rare long distance dispersal (< 1,500 km). Theory predicts that gene flow reduces local adaptation [105], and small populations governed by strong genetic drift (e.g. Madeira) are less likely to have fine-scale local adaptations [106]. Therefore, translocation among inbred nearby populations presents limited risks of outbreeding depression [107,108]. The success of such genetic rescue is however conditioned by long-term conservation of extant populations, their habitat and resources, and actions to re-establishing natural dispersion among populations [108].

Finally, our broad scale genetic survey provides a new turnkey cost-efficient tool to accurately trace the origin of vagrant MMS individuals and monitor the genetic diversity of MMS populations and colonies. The combination of the 19 MMS microsatellites markers with the CR1 mitochondrial sequence may indeed serve not only to select the MMS individuals for potential translocation, but also to assess translocation success over time [104].

## Data accessibility

Microsatellite data is available in table S1. All additional data, scripts and materials are available to readers at https://doi.org/10.5281/zenodo.6871982.

## Acknowledgments

We thank all the collectors and museums listed in Table S2 for providing access to genetic samples. We are grateful to Sophie Courjal and the staff of the “Plateau technique - Biologie moléculaire et microbiologie” at EDB for their assistance during lab work, to P. Kyritsis [Archipelagos], for his logistic help, to I. Carvalho for early comments on the manuscript and two anonymous reviewers who significantly helped to improve the manuscript. This work was funded by the Fondation Prince Albert II de Monaco (project “Génétique de la conservation du phoque moine de Méditerranée”). The Genotoul bioinformatics (Bioinfo Genotoul) platforms provided computing resources. JS was supported by PANGO-GO (ANR-17-CE02-0001), and LABEX TULIP (ANR-10-LABX-0041).

## Conflict of interest disclosure

The authors of this article declare that they have no financial conflict of interest with the content of this article.

## Authors’ contribution

PG, EL and JS designed the study. AA, AAK, PFL, RP, GM, AP, TA, SA, AB, ED, BO, AMT, LMG and PD collected the genetic samples and supplied curated databases. JD and PG did the lab work. JS conducted data analyses. JS and PG drafted a first version of the manuscript. All co-authors participated in the writing of the manuscript and agreed with its last version.

## Notes

https://doi.org/10.5281/zenodo.6871982

### Competing Interest Statement

The authors have declared no competing interest.

### Summary of Updates

This version of the manuscript has been revised according to two rounds of review by two reviewers' comments and suggestions during a submission process at Proceedings of the Royal Society B.

