## Supporting Information for "The antique genetic plight of the Mediterranean monk seal (*Monachus monachus*)"

doi:10.1098/rsbpb.2022-0846

Supporting information for: The antique genetic plight of the Mediterranean monk seal (*Monachus monachus*). J. Salmona et al., in revision. doi:10.1098/rspb.2022-0846.

#### Table of content:

**Supporting Methods** 4
Method S1: DNA Extraction 4
Method S2: PCR Amplification of mtDNA Control Region 5
Method S3: Microsatellites genotyping 5
Genotyping 5
Sample, missing data filtering and allelic dropout 6
Markers quality control 7
Method S4: Genetic structure analyses 7
Method S6: Approximate Bayesian Computation 9
Mutation rate and generation time 9
Models 9
Missing data, microsatellites and mtCR1 integration 10
Summary statistics 10
Partial least square regression 10
Minimum entropy 10
Model selection 10
ABC validation and parameter estimation 11

**Supporting Tables** 12
Table S1. Sampling summary 12
Table S2. Samples and genotypes information 12
Table S3. mtDNA diversity and haplotype frequencies 13
Table S4. Microsatellite null alleles across populations 13
Table S5. Custom migration matrix among MMS populations 14
Table S6. ABC model parameters and priors 14
Table S7. ABC observed summary statistics for modern samples 14
Table S8. ABC observed summary statistics for modern and historic populations 15
Table S9: ABC model fit 15
Table S10: ABC structured model fit with selected statistics 16
Table S11: ABC  parameter posterior estimates 16

**Supporting Figures** 17
Figure S1. Map of the Mediterranean monk seals included in the study 17
Figure S2. Detailed maps of the MMS samples included in the study 18
Figure S3. MMS sampling distribution through time 19
Figure S4. Microsatellites linkage disequilibrium across all samples 20
Figure S5. Microsatellites linkage disequilibrium WS samples 21
Figure S6. Microsatellites linkage disequilibrium EM samples 22
Figure S7. Microsatellites Hardy-Weinberg equilibrium 23

Figure S8. Microsatellites genotype accumulation curve 24
Figure S9: Summary of demographic models tested using ABC. 25
Figure S10: ABC panmictic model choice 26
Figure S11: ABC structure choice 27
Figure S12: ABC structured stationary model choice 28
Figure S13: ABC complex model choice 29
Figure S14: ABC final model choice 30
Figure S15: Most likely number of panmictic clusters in MMS 31
Figure S16: MMS genetic clustering 32
Figure S17: DAPC representation of microsatellites structure 33
Figure S18: PCA representation of microsatellites structure. 34
Figure S19. Monachus monachus mitochondrial genetic diversity. 35
Figure S20. Temporal evolution of MMS genetic diversity. 36
Figure S21: Temporal evolution of the Western-Saharan/Mauritania genetic diversity 37
Figure S22: Temporal evolution of the east-Med genetic diversity 38
Figure S23: MMS d-loop haplotypes distribution over time 39
Figure S24: MMS d-loop haplotypes geographic repartition over time 40
Figure S25: Western-Sahara/Mauritania 1997 die-off effect on genetic diversity 41
Figure S26: Western-Sahara/Mauritania sampling temporality 42
Figure S27: Cabo Blanco 1997 die-off effect on rare allele frequencies 43
Figure S28: Cabo Blanco 1997 die-off effect on major allele frequencies 44
Figure S29: Geographic scale influence on IBD 45
Figure S30: Isolation by distance in MMS 46
Figure S31: ABC population size posteriors 47
Figure S32: ABC migration rate posteriors 48
Figure S33: Bayes factor of period of MMS' decline. 49

**Supporting References** 50

### Supporting Methods

#### Method S1: Sampling and DNA Extraction

The MOm sample collection was conducted in accordance with the guidelines of the relevant research permits (86286/340, 184316/4337, 165523/88, 118956/3033, 151241/138, 178104/66, and 1893/43) issued by the Hellenic Ministry of Environment and Energy. The University of Barcelona samples were collected in the 1990s and followed to good practices issued by: i) the European Union funding agency which specially set-up a Committee of Experts that supervised the fieldwork and all sample collection; ii) the project was co-conducted by the Spanish Directorate for the Conservation of Nature of the Minisitry of the Environment, and in most cases officials were personally involved in the collection and transport of samples. The Archipelagos samples were exclusively hair samples collected between 1991 and 1999 in monk seal caves in the Ionian Sea, Greece, after molting and not affecting animals. Finally, the Turkish routinely collected necropsies samples were sent under the CITES export permit TR1200215 - - 044, issued by the Turkish Ministry of Food Agriculture and Livestock, General Directorate of Fisheries and Aquaculture.

Genomic DNA was extracted from fresh tissue and skin samples with the NucleoSpin Tissue kit (Macherey-Nagel, Hoerdt, France), following manufacturer's recommendations. Final elution step was repeated twice using 50 + 50 µl Elution buffer to increase DNA yield.

To avoid cross-sample contamination, bones (postcranial and turbinal bones), hairs, feces, tanned skins and connective tissues were processed by series of small batches (ca. 10 samples each) with negative controls through each extraction step in a room dedicated to degraded and environmental DNA (DNA Trace, EDB - Toulouse), isolated from the other lab facilities where fresh tissues were processed. A subset of 2x10 samples was extracted 2 – 3 times independently and sequenced to control for potential cross-contamination and mismatch repair.

Hairs and samples of tanned skin and connective tissues were extracted using a modified CTAB protocol (Gaubert & Zenatello, 2009) including upstream TE washing baths and the addition of 4 µL DTT (1M) and 20 µL proteinase K (20mg/mL) during the lysis step. Final elution volumes in nuclease free water varied from 25 to 120 µl, depending on the size of the DNA pellet.

Bones were reduced to powder using a Bel-Art mortar (Fisher Scientific, Illkirch-Graffenstaden, France) cooled with liquid nitrogen. Approximately 60 – 140 mg of bone powder was digested in Buffer A (for 100 samples: 5mL Tris 1M, 2mL EDTA 0.5M, 2.5mL N-Lauryl Sarcosyl 20%, 3.55g Sodium hydrogen phosphate, 1mL Proteinase K 40U/mg) during 15minutes at 37°C, then in Buffer B (for 100 samples: 100mL EDTA 0.5M, 3.55g Sodium hydrogen phosphate, 200uL Proteinase K 40U/mg) during 48h of incubation with soft shaking at 37°C. Final elution was done with EBT Buffer (20mL of EB Buffer and 10 uL of Tween20 0.05%) in a volume of 60uL. Samples with original quantity of material <60 mg were processed with half the volumes indicated above.

Feces were rehydrated in 500uL of DNA free water with hand shaking during 2 minutes, then reduced in powder. Fecal DNA was subsequently processed with Macherey-Nagel™ NucleoSpin™ DNA Stool columns following the manufacturer's guidelines.

DNA concentrations were estimated on a NanoDrop 1000 Spectrophotometer (ThermoFisher Scientific, Illkirch-Graffenstaden, France).

#### Method S2: PCR Amplification of mtDNA Control Region

We amplified 524 bp of the hypervariable region I of the mitochondrial control region (CR1) from the fresh and hair samples after (Karamanlidis *et al.*, 2016). For the historical samples, we used the four specific primer pairs designed by (Gaubert *et al.*, 2019) to obtain a 484 bp fragment encompassing all the variable sites of CR1 observed in MMS. In this case, negative DNA-extraction controls (see above) were included along with PCR blank controls to further assess potential contamination. PCR amplifications were carried out for both the large and small fragments in 20 µL final volume containing ~5 – 50 ng of template DNA, 0.1 mg/mL BSA, 4 × 0.25 mM dNTPs, 2 × 0.2 mM primers, 1 × PCR direct loading buffer with MgCl 2 (1.5 mM) and 0.5 U GoTaq Flexi DNA Polymerase (Promega, Charbonnières-les-Bains, France). PCR cycling conditions included a first step of denaturation (94°C, 2 min), followed by 40 – 45 cycles of denaturation (92°C, 30 s), annealing (30s) and extension (72°C, 30 s), and a final extension step (72°C, 15 min). PCR products were sequenced in both directions on a AB3130xl Genetic Analyzer (Applied Biosystems, Foster City, CA) at Genoscreen (Lille, France). We completed our data set with 165 publicly available sequences [GenBank accession numbers: MG570469-570475 and KT935307-935311, (Karamanlidis *et al.*, 2016; Gaubert *et al.*, 2019) respectively] covering the extant and historical species' range. Nucleotide sequences were visually aligned with BioEdit 7.1.3 (Hall, 1999). The final CR1 alignment included 328 sequences (Table S1).

#### Method S3: Microsatellites genotyping

##### Genotyping

We genotyped 383 samples at 19 nuclear microsatellite loci ((Dayon *et al.*, 2020), Method S3) from 314 modern samples and 69 historical samples, using the procedure described in Dayon et al., 2020. We used four PCR multiplexes (4 – 5 loci) with four different ABI fluorescent dyes (Dayon *et al.*, 2020). PCR amplifications for each multiplex were carried out in a total 20 µl reaction mixture containing approximately 20 ng of genomic DNA, 1x Multiplex PCR Master Mix (QIAGEN Multiplex PCR Plus Kit; Qiagen, Courtaboeuf, France) and 0.2 µM of each primer pair. PCR thermoprofiles included an initial denaturation step (95 °C for 5 min), followed by 32 cycles of denaturation (95 °C for 30 s) — annealing (60 °C for 90 s) — elongation (72 °C for 30 s), and a final elongation step (60 °C for 30 min). To improve the scoring of alleles, we proceeded to random, repeated (× 2-5) amplifications of 1-to-3 multiplexes in ca. 75% of the samples (Dayon *et al.*, 2020). PCRs of DNA extracts from hairs, feces, and museum material were systematically replicated 2-to-5 times. In order to restrain allelic dropout, alleles were only scored when a majority consensus was reached among PCR replicates for a given locus/sample. The PCR products were run on an ABI 3730 DNA Analyzer (Thermo Fisher Scientific) at Genoscreen (Lille,France; https:// www.genoscreen.fr). Allele scoring and final extraction of genotypes were performed in Geneious v 9.0.5 (Kearse *et al.*, 2012) with the Microsatellites plugin (https://www.geneious.com/features/microsatellite-genotyping/).

#### Sample, missing data filtering and allelic dropout

Out of the 383 starting samples, 23 (20 historical, 3 modern) simply did not work at all, and 37 (20 historical, 17 modern) did successfully amplify at one locus only (out of 19 loci). We then applied a two-fold sample selection procedure to optimize the number of individuals genotyped at informative loci. We relied on the minimum number of loci necessary to discriminate among individuals and the region-based discriminant analysis of principal components [DAPC] contribution of loci. This procedure, overcoming the use of simple threshold not considering loci informativeness, allowed filtering-out individuals genotyped mostly for little informative loci, while keeping individuals genotyped mostly for few but informative markers. Using a selection of 277 samples genotyped for at least 65 % of the loci, we estimated to 10 the minimum number of loci necessary to discriminate between individuals (Fig. S8) using the R package *poppr* (Kamvar *et al.*, 2014). Additionally, using the R package *Adegenet* (Jombart, 2008) we extracted the loci contributions to the first two components of a DAPC analysis considering main sampling regions as prior groups (Western-Sahara, Madeira, Western-Mediterranean, Central-Mediterranean, Eastern-Mediterranean). This approach allowed identifying the loci that carry most geographically relevant information revealing the genetic structure of the MMS. We further filtered the initial dataset, applying a weighted score to each locus based on their contributions to the first two components of the abovementioned DAPC analysis. This procedure allowed filtering out individuals successfully genotyped for mostly little informative loci, while keeping individuals genotyped for fewer but more informative markers. Using this procedure, 44 samples (15 historical, 29 modern) that amplified successfully in only 2-6 loci were not retained. Above this threshold (7 loci amplified at least), 14 samples (13 modern and 1 historical) were discarded. These filters reflect the degree of missing locus and of informativeness of the dataset, and are not indicative of any putative allelic dropout. As a token of transparency, we had chosen to expose the full numbers of samples that were DNA extracted for the study, and to explain the quality / informativeness filters that we have used to build up a solid dataset.

We further assessed the presence of potential duplicates in our data using the *psex*, *mlg and mlg.filter* functions of the R package *poppr* and considering missing data (https://doi.org/10.5281/zenodo.6871982). Potential duplicates were confronted to their available information in the database (provenance, age at sampling, sex, collection, ID number similarities). Conflicting cases (e.g. “Female 1985 from WS” and a “Male pup 2004 from WS”) were considered as identical genotypes, while one of the two identical genotypes was removed in the non-conflictual cases. Following that procedure, we removed 10 potential modern duplicates.

Altogether, a total of 56 (/70) historical and 72 (/313) modern samples did not pass our sequential filtering procedure. All historical, fecal, and hair samples were PCR-replicated 2-to-5 times (as indicated in the manuscript), with museum samples being replicated at least 3 times, to ensure genotype accuracy and to limit potential allelic dropout. A number of simple indices suggest that the historical samples that we used in our analyses were not affected by allelic dropout (or were similarly affected compared to fresh samples). The 14 historical and 241 modern retained samples showed similar levels of missing data (median: 0.083 & 0.087; sd: 0.27 & 0.07 respectively, Tukey-HSD p-value = 0.15). The tests conducted in Microchecker, per population, and across the whole data, did not detect evidence of large allele dropout, for any of the genotyped loci (Table S3). The individual levels of historical samples inbreeding were lower or exceptionally within the range of those estimated for modern individuals (Fig. 2e).

##### Markers quality control

None of the marker pairs were consistently in significant genotypic linkage disequilibrium across the three tested dataset (all 253 samples [Fig. S4], 96 Western-Sahara [Fig. S5], and 148 eastern-Mediteranean [Fig. S6] specimens respectively) when tested using the Agapov and Burt $\mathbf{r_d}$ linkage disequilibrium (Agapow & Burt, 2001) in the R package *poppr* (Kamvar *et al.*, 2014, 2015). Five loci (MM_395180; MM_25313; MM139103; MM_821171; MM_989657) showed evidence of null alleles in MICRO-CHECKER (Van Oosterhout *et al.*, 2004) across the two major tested populations (W-Sahara & Eastern-Mediteranean; Table Sx null alleles MC). One locus (MM_821171) departed significantly from Hardy – Weinberg equilibrium (Fig. Sx_HWEtest) as estimated using Nei $F_{IS}$ statistics (Nei, 1977) and a $\chi^2$ exact test based on 1,000 Monte Carlo permutations using the R packages *demerelate* (Kraemer & Gerlach, 2017) and *pegas* (Paradis, 2010), respectively. We did not remove these loci, as homozygote excess and departure from Hardy – Weinberg equilibrium at a locus can result from genotyping errors as well as from the genetic characteristics of populations [Inbreeding; structure; (Wittke-Thompson *et al.*, 2005; Campagne *et al.*, 2012)].

The final microsatellites data set comprised 19 loci genotyped for 253 samples (including 12 historical samples; Table S1) with 10.18 % missing data.

#### Method S4: Genetic structure analyses

**Data structure**

The relative complexity of the dataset (nuclear microsatellites and mtDNA sequences / modern and historic samples) lead to multiple analyses involving distinct subsets of data. While several "naive" analyses were conducted with all samples such as structure, assignment and haplotype network, others included only a category of samples (modern vs. historic). Further analyses included a selection of these when focusing on a particular locality (e.g. modern Western-sahara/Mauritania pre/post die-off). Data set compositions are detailed below ad-hoc figures and tables, and their construction is reproducible with the scripts available at https://doi.org/10.5281/zenodo.6871982.

**Genetic diversity**

To assess the overall, per sampling site, and per locus microsatellites genetic diversity, we estimated the allelic richness ($A_R$ (Hurlbert, 1971)) using minimum sampling sizes of 5, 10, and 25 for rarefaction with the R package *hierfstat* (Goudet, 2005). We also assessed the number of alleles (*A*), the observed ($H_O$) and Nei's unbiased expected heterozygosities ($H_E$ (Nei, 1978)) with the R package *adegenet* (Jombart, 2008). These metrics were estimated per sampled basin or per cluster, using either modern [all sampled basin] or historical samples [Western-Mediterranean samples]. Additional analyses also compared overall estimates [all historical *vs* all modern samples]. Analyses of the 1997 die-off in Cabo Blanco do compare modern individuals before and after the 1997 die-off. Pre-die-off samples include all individuals from 1975 to 1997. Post-die-off samples include only individuals that were born after 1997. To evaluate the effect of drift in the small colonies of MMS, we estimated the average individual inbreeding coefficient *F* (Jombart, 2008) in *adegenet*, from 100 iterations.

**Genetic structure**

We first assessed the level of microsatellites genetic differentiation among localities, populations, and time periods through Nei's $F_{ST}$ (Nei, 1973) with the R package *diveRsity* (Keenan *et al.*, 2013). Additionally, in *adegenet*, we investigated patterns of genetic variance with principal component analysis (PCA) of allele frequencies and, with a discriminant analysis of principal components (DAPC) considering sampling regions as groups. We further assessed the structure using the SnapClust clustering approach (Tonkin-Hill *et al.*, 2019) with default values for $K$ = 1-15, and selected the most appropriate $K$ value using the goodness of fit Akaike, Bayesian and Kullback information criterion (Akaike, 1974; Cavanaugh, 1999; Akogul & Erisoglu, 2016). MtDNA haplotype relationships were reconstructed with a maximum parsimony network using *pegas* with the default parameters.

**Analyses of connectivity**

To assess the potential effect of distance on gene flow, we investigated patterns of isolation by distance (IBD) through Mantel tests (Mantel, 1967) with 999 permutations, using the R package *ade4* (Chessel *et al.*, 2004). Individual geographical distances and Euclidean, Bruvo's (Bruvo *et al.*, 2004), Prevosti's (Prevosti *et al.*, 1975), and Kosman's (Kosman & Leonard, 2005) genetic distances were estimated and compared using the R packages *poppr* (Kamvar *et al.*, 2014) and *PopGenReport* (Adamack & Gruber, 2014). Since IBD may be limited to a certain scale (Keller & Holderegger, 2013; Van Strien *et al.*, 2015), we compared subsets of pairwise data defined by a maximum geographic distance (*S*) between samples (Cayuela *et al.*, 2019). *S* ranges from 50 km [the estimated distance at which no MMS individual is excluded from a neighboring graph (Jombart, 2008)] to 6,000 km (the maximum Euclidean distance between two individuals in our study), by 100 km increments. For each subset, we ran the IBD test and plotted the model fit ($R^2$) against *S* to assess which spatial scale optimizes the amount of variance explained (Cayuela *et al.*, 2019). In addition, we computed Mantel correlograms (Oden & Sokal, 1986; Sokal, 1986) using the R package *vegan* (Dixon, 2003), with 50 classes of ~100km width, and a multiple-testing corrected assessment of Mantel statistics through 999 permutations. We used Nei's $F_{ST}$ (Nei, 1973) for population-based inferences of IBD.

### Method S6: Approximate Bayesian Computation

Traditional approaches reconstructing demographic history assume that samples are obtained from isolated populations, hence ignoring genetic substructure and migration from other populations. A growing number of studies showed that ignoring population substructure or using an inadequate sampling scheme may lead to spurious signatures of demographic change (Leblois *et al.*, 2006; Chikhi *et al.*, 2010; Heller *et al.*, 2013; Mazet *et al.*, 2015, 2016). We reconstructed the demographic history of the Mediterranean monk seal using ABC approaches implemented in ABCTOOLBOX version 2.0 (Wegmann *et al.*, 2010) and in the R package *ABC* (Csilléry *et al.*, 2012). The principle behind ABC is to compare data simulated under several alternative scenarios to the real data, using (in general) summary statistics. Alternative scenarios can subsequently be compared and parameters of interest estimated from the most supported scenarios (Csilléry *et al.*, 2010). Within this framework, we simulated genetic data using the coalescent tool FASTSIMCOAL version 2.6.0.3 (Excoffier & Foll, 2011).

#### Mutation rate and generation time

Mutation rate (Stoffel *et al.*, 2018) and generation time (Karamanlidis & Dendrinos, 2015) were extracted from existing literature, in line with the range of empirical estimates (Ellegren, 2004; Selkoe & Toonen, 2006).

#### Models

We first tested scenarios assuming a single panmictic population and different histories of population size change (Figs S9-10). These models were tested using modern samples, because separating historic populations from modern ones would require including population structure in the models.

Second, we modeled stable-size structured populations with constant migration rates among all populations or among the closest geographically, respectively using the *n-island* or stepping-stone models (Figs S9, S11). These models additionally included (i) 5-10 unsampled ghost populations mimicking extinct populations and (ii) a change in connectivity at $t_1$, mimicking the loss of gene flow among extant populations (Figs S9, S11-12).

Third, we combined population size change, structure, ghost populations, and changes in connectivity, to model populations that suffered one or several events of decline and fragmentation (Figs S9, S13). These scenarios were constructed using the stepping-stone model, or spatially-explicit schematization of the extinct and extant populations allowing to attribute a particular gene flow class to each pair of populations based on their distance and position in space (Table S4). These last two sets of models (second and third) were tested using modern data only, and modern data along with the historical Western-Mediterranean population sampled ~140 years in the past. All simulations assumed log-uniform priors for population sizes, migration rates and time of demographic events (Table S5).

#### Missing data, microsatellites and mtCR1 integration

The mtDNA d-loop data set was reduced to individuals common with the microsatellites data set (N = 199). Other individuals were assigned missing data for the mtDNA d-loop loci. To include both microsatellites and mtDNA within the ABCTOOLBOX analytical frame, we used a home made bash script that splits the two types of markers from each simulation prior to the statistics estimations, and joins the statistics afterwards. Prior to summary statistics estimation, we also replicated the missing data pattern of the observed data onto the simulated data-set using custom awk scripts (https://doi.org/10.5281/zenodo.6871982).

#### Summary statistics

We used ARLSUMSTAT (Excoffier & Lischer, 2010) to estimate summary statistics from observed and simulated data (Tables Sx_ABCssmod & Sx_ABCssall). To overcome the potential overfitting of parameters, we reduced the summary statistics dimension (Blum *et al.*, 2013), both using partial least square regression (PLS) regression and minimizing the sample entropy (Nunes & Balding, 2010).

##### Partial least square regression

To reduce the high dimensionality of structured scenario's summary statistic data we applied a partial least square (PLS) regression using the R script provided in the ABCTOOLBOX package. We chose the optimal set of PLS components (i.e. the smallest set carrying a large amount of information about the model parameters) by assessing the root-mean-squared error for each parameter. This allowed us to reduce a large set of summary statistics to a smaller number of independent components, in which informative summary statistics are weighted more than those that do not respond to changes in parameter values (Wegmann *et al.*, 2009).

##### Minimum entropy

We additionally inferred the most informative summary statistics subset, by minimizing the sample entropy as a proxy of a posterior sample measure, across all possible subsets of summary statistics (Nunes & Balding, 2010), in abctools (Nunes & Prangle, 2015). For computational reasons, we run five sequential estimates of the most informative summary statistics subset, using the results of the precedent analysis as input. Across models, we retained the final summary statistic subset, best improving the model choice statistics, and therefore exhibiting a low MD, high proportions (MD *p-value*) of retained simulations showing a lower or equal likelihood under the inferred GLM as compared to the observed genetic data (Wegmann *et al.*, 2010), and with high centrality of the observed data within the multidimensional cloud of retained simulations [Tukey *p-value* (Kousathanas *et al.*, 2018)]. The complete procedure can be found at https://doi.org/10.5281/zenodo.6871982.

#### Model selection

To assess model fit, we first calculated the marginal densities (MD) with the generalized linear model (GLM) built from the 10,000-1,000 simulations (i.e., 0.2-0.02% of the 5x$10^5$) closest to the observed data in ABCTOOLBOX. We retained models with low MD, high proportions (MD *p-value*) of retained simulations showing a lower or equal likelihood under the inferred GLM as compared to the observed genetic data (Wegmann *et al.*, 2010), and with high centrality of the observed data within the multidimensional cloud of retained simulations [Tukey *p-value* (Kousathanas *et al.*, 2018)]. To compare models we estimated the Bayes factor (BF), the ratio of the posterior densities of alternative hypotheses (i.e., scenario), over the ratio of their prior densities. BF absolute values > 3 were considered as significant evidence to reject the alternative hypothesis (Kass & Raftery, 1995). When two models showed BF absolute values < 3, we kept the simplest model but considered both models' results for discussion.

#### ABC validation and parameter estimation

We first investigated ABC's ability to distinguish between the proposed models, with 1000 pods randomly selected from simulated data sets under each model. Following the same assignment procedure as for the observed data, we derived the model misclassification rate by counting all pods assigned to a model other than the one generating it, and summarized them in a so-called confusion matrix (Csilléry *et al.*, 2012).

We assessed the accuracy of parameter moments estimates (mode, mean and median) under several tolerance levels (0.05; 0.01; 0.005; 0.001; 5e-04; 1e-04; proportion of retained simulations, i.e. 50,000 to 100 simulations) with the leave-one-out cross-validation procedure (with 50 simulations randomly selected as pods) available in the R package "abc" (Csilléry et al., 2012). The summed difference between the pods and estimated parameters were first used to calculate the precision of each parameter estimate (equation 4 in Csilléry et al. 2012). A prediction error of >1 suggests an uninformative model or simply that the prior is equal to the posterior: values closer to zero suggest the parameter of interest can be reliably estimated with the model. We also estimated the average of the root-mean-squared errors [RMSE; (Leuenberger & Wegmann, 2010)] over the 50 pods for the 6 tolerance levels.

To increase the accuracy of posterior distribution estimates from the best-fitting models, we produced $5x10^6$ simulated data sets under these models and estimated posteriors using the GLM approach with 0.01% (i.e., 500) simulations closest to the observed data.

To compare alternative temporally delineated hypotheses and identify the most likely time of demographic events we performed a BF analysis. We identified seven time intervals corresponding to putative causes of MMS demographic events.

### Supporting Tables

#### Table S1. Sampling summary

Table presenting the number of samples genotyped and retained across methods and localities. Epoch: His: historical samples, Mo: modern samples, Un: samples of unclear date, Ov: Overall (sum). Intersect A|B: samples genotyped for both the 19 microsatellites loci and at the mtDNA locus CR1.

| Data | W-Sahara | | Madeira | | W-Med | | | C-Med | | | E-Med | | | Black-Sea | | Total | | | |
|---|---|---|---|---|---|---|---|---|---|---|---|---|---|---|---|---|---|---|---|
| Epoch | His | Mo | His | Mo | His | Mo | Un | His | Mo | Un | His | Mo | Un | His | Mo | His | Mo | Un | Ov |
| Microsatellites tested | 4 | 141 | 4 | 11 | 40 | 2 | 0 | 11 | 29 | 0 | 10 | 130 | 0 | 1 | 0 | 70 | 313 | 0 | 383 |
| Microsatellites retained (A) | 3 | 93 | 0 | 10 | 7 | 2 | 0 | 3 | 26 | 0 | 0 | 108 | 0 | 1 | 0 | 14 | 239 | 0 | 253 |
| mtDNA (B) | 2 | 134 | 7 | 14 | 26 | 2 | 1 | 3 | 12 | 1 | 0 | 142 | 2 | 4 | 0 | 42 | 304 | 4 | 350 |
| Intersect A\|B | 1 | 66 | 0 | 10 | 7 | 2 | 0 | 2 | 9 | 0 | 0 | 108 | 0 | 0 | 0 | 10 | 195 | 0 | 205 |
| All samples (A&B) | 4 | 161 | 7 | 14 | 26 | 2 | 1 | 4 | 29 | 1 | 0 | 142 | 2 | 5 | 0 | 46 | 348 | 4 | 398 |

#### Table S2. Samples and genotypes information

Table presenting the samples used in the study, their metadata and microsatellite genotypes. ID: sample identifier (PG database); Region: sampling region, Simplified_region: larger area than the latter for which regions were grouped together to form larger entities on which population genetics statistics can be estimated; Longitude & Latitude: approximate sample geographic coordinates (in WGS84-degree decimal); Year: sampling year as reported by collectors; Simplified_year: approximated sampling time only modified for non-integer values; Period: categorized period of sampling modern (≧ 1980), historic (< 1980); Country: sampling country; Island locality: more precise name of the sampling locality when available; Age: age of the animal at sampling; Sex: sex of the animal; NS: specimen not sequenced at the mtDNA CR1 locus; #called_alleles: number of called alleles; #uncalled_alleles: number of uncalled alleles; Missing rate: sample based loci missing rate.

#### Table S3. mtDNA diversity and haplotype frequencies

Table presenting the haplotypes and the summary statistics inferred from the 328 Mediterranean monk seals CR1-mtDNA sequence data, for each sampled region. Lit. ID: Correspondence with Haplotypes identifiers used in the literature (Karamanlidis *et al.*, 2016; Gaubert *et al.*, 2019; Rey-Iglesia *et al.*, 2021); Hap. ID; new identifiers as obtained from the analyses (https://doi.org/10.5281/zenodo.6871982); Total: sum of the number of haplotypes (does not repeat C-E-Med), *h*: haplotype diversity (Nei & Tajima, 1981), and nucleotide diversity *π* (Nei, 1987), W-Sahara: Western-Sahara/Mauritania, W-Med: Western Mediterranean Sea, C-Med: Central Mediterranean Sea, E-Med: Eastern Mediterranean Sea, C-E-Med: Central and Eastern Mediterranean Sea considered together, Unknown: sample with unclear origin.

| | **Haplotypes** | | | | | | | **Statistics** | | | |
|---|---|---|---|---|---|---|---|---|---|---|---|
| **Lit. ID** | **MM01** | **MM02** | **MM03** | **MM04** | **MM05** | **MM06** | **MM07** | | | | |
| **Hap. ID** | **I** | **II** | **IV** | **V** | **III** | **VI** | **VII** | **Total** | $n_h$ | $h$ | $\pi$ |
| W-Sahara | 1 | 0 | 0 | 0 | 134 | 1 | 0 | 136 | 3 | 0.029 | 0.00006 |
| Madeira | 0 | 0 | 0 | 0 | 20 | 0 | 0 | 20 | 1 | 0.000 | 0.00000 |
| W-Med | 14 | 0 | 0 | 3 | 7 | 0 | 4 | 28 | 4 | 0.680 | 0.00272 |
| C-Med | 3 | 0 | 12 | 0 | 0 | 0 | 1 | 16 | 3 | 0.425 | 0.00145 |
| E-Med | 110 | 36 | 0 | 1 | 0 | 0 | 0 | 147 | 3 | 0.383 | 0.00083 |
| **C-E-Med** | 113 | 36 | 12 | 1 | 0 | 0 | 1 | 163 | 5 | 0.468 | 0.00114 |
| Black-Sea | 0 | 4 | 0 | 0 | 0 | 0 | 0 | 4 | 1 | 0.000 | 0.00000 |
| Unknown | 0 | 0 | 1 | 0 | 0 | 0 | 0 | 1 | 1 | NA | NA |
| Total | 128 | 40 | 12 | 4 | 161 | 1 | 5 | 351 | 7 | 0.643 | 0.00203 |

#### Table S4. Microsatellite null alleles across populations

Table presenting the comparison of estimated null allele frequencies analyses conducted in MICRO-CHECKER (Van Oosterhout *et al.*, 2004). Null: final evidence for null allele presence per data set as returned by MICRO-CHECKER; The estimated null allele frequency for each locus is compared to the null allele frequencies obtained using methods by Chakraborty (Chakraborty *et al.*, 1992) and Brookfield (Brookfield, 1996). Homozygote excess: analysis indicates homozygote excess at this locus; Large allele dropout: analysis indicates that large alleles do not amplify as efficiently as small alleles at this locus.

#### Table S5. Custom migration matrix among MMS populations

Spatially-explicit schematization of the extinct and extant populations (left) used to attribute a particular gene flow class to each pair of populations (right) based on their simplified distance and position in space, for 11 populations (upper panel) and 12 populations (lower panel). Gene flow classes are subsequently used in ABC modeling. Cells with value 1 exchange individuals at the rate sampled from the prior distribution. Cells with value 2 exchange ⅓ of cells with 1, and those with 3 exchange ⅓ of cells with 2.

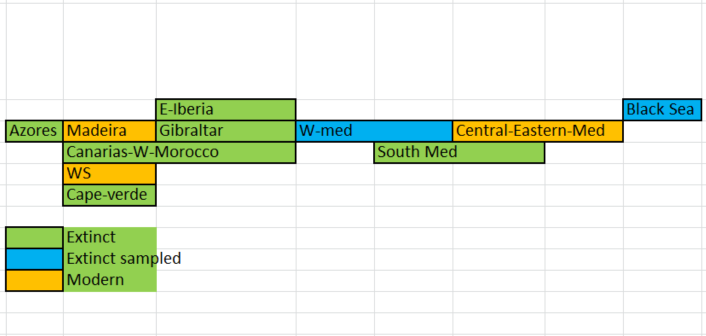


**Intuitive matrix**

| | | | Black-Sea | CE-Med | S-Med | W-med | Gibraltar | E-Iberia | Madeira | Azores | Canarias | WS | Cape-verde |
|---|---|---|---|---|---|---|---|---|---|---|---|---|---|
| Ghost | 1 | Black-Sea | 0 | 1 | 2 | 3 | 0 | 0 | 0 | 0 | 0 | 0 | 0 |
| Sampled | 2 | CE-Med | 1 | 0 | 1 | 1 | 3 | 0 | 0 | 0 | 0 | 0 | 0 |
| Ghost | 3 | S-Med | 2 | 1 | 0 | 1 | 2 | 3 | 0 | 0 | 3 | 0 | 0 |
| Sampled | 4 | W-med | 3 | 1 | 1 | 0 | 1 | 2 | 2 | 3 | 2 | 3 | 0 |
| Ghost | 5 | Gibraltar | 0 | 3 | 2 | 1 | 0 | 1 | 1 | 2 | 1 | 2 | 3 |
| Ghost | 6 | E-Iberia | 0 | 0 | 3 | 2 | 1 | 0 | 1 | 1 | 2 | 3 | 0 |
| Sampled | 7 | Madeira | 0 | 0 | 0 | 2 | 1 | 1 | 0 | 1 | 1 | 2 | 3 |
| Ghost | 8 | Azores | 0 | 0 | 0 | 3 | 2 | 1 | 1 | 0 | 2 | 3 | 2 |
| Ghost | 9 | Canarias | 0 | 0 | 3 | 2 | 1 | 2 | 1 | 2 | 0 | 1 | 2 |
| Sampled | 10 | WS | 0 | 0 | 0 | 3 | 2 | 3 | 2 | 3 | 1 | 0 | 1 |
| Ghost | 11 | Cape-verde | 0 | 0 | 0 | 0 | 3 | 0 | 3 | 2 | 2 | 1 | 0 |

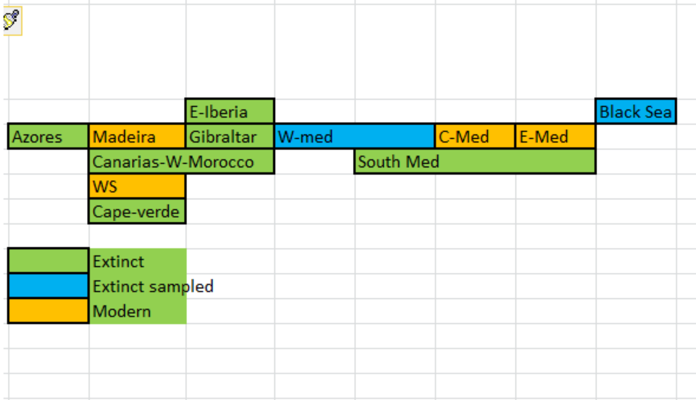


**Intuitive matrix**

| | | | Black-Sea | CE-Med | C-Med | S-Med | W-med | Gibraltar | E-Iberia | Madeira | Azores | Canarias | WS | Cape-verde |
|---|---|---|---|---|---|---|---|---|---|---|---|---|---|---|
| Ghost | 1 | Black-Sea | 0 | 1 | 2 | 2 | 3 | 0 | 0 | 0 | 0 | 0 | 0 | 0 |
| Sampled | 2 | E-Med | 1 | 0 | 1 | 1 | 2 | 3 | 0 | 0 | 0 | 0 | 0 | 0 |
| Sampled | 3 | C-Med | 2 | 1 | 0 | 1 | 1 | 2 | 3 | 0 | 0 | 0 | 0 | 0 |
| Ghost | 4 | S-Med | 2 | 1 | 1 | 0 | 1 | 2 | 3 | 0 | 0 | 3 | 0 | 0 |
| Sampled | 5 | W-med | 3 | 2 | 1 | 1 | 0 | 1 | 2 | 2 | 3 | 2 | 3 | 0 |
| Ghost | 6 | Gibraltar | 0 | 3 | 2 | 2 | 1 | 0 | 1 | 1 | 2 | 1 | 2 | 3 |
| Ghost | 7 | E-Iberia | 0 | 0 | 3 | 3 | 2 | 1 | 0 | 1 | 1 | 2 | 3 | 0 |
| Sampled | 8 | Madeira | 0 | 0 | 0 | 0 | 2 | 1 | 1 | 0 | 1 | 1 | 2 | 3 |
| Ghost | 9 | Azores | 0 | 0 | 0 | 0 | 3 | 2 | 1 | 1 | 0 | 2 | 3 | 2 |
| Ghost | 10 | Canarias | 0 | 0 | 0 | 3 | 2 | 1 | 2 | 1 | 2 | 0 | 1 | 2 |
| Sampled | 11 | WS | 0 | 0 | 0 | 0 | 3 | 2 | 3 | 2 | 3 | 1 | 0 | 1 |
| Ghost | 12 | Cape-verd | 0 | 0 | 0 | 0 | 0 | 3 | 0 | 3 | 2 | 2 | 1 | 0 |

#### Table S6. ABC model parameters and priors

Details of ABC models parameters and priors. Nb_pop: number of populations, obs_file, ssdef, ref_file: .obs, .ssdef and .arp files used in ABCTOOLBOX, Gen_time: generation time used to draw priors, T0-T1-T2: demographic event time priors, N0-N1-N2: effective population size priors, NM0-NM1: migration rate priors, prior_min-max: log10 scaled prior values, prior_years-diploid-diploi/gen: natural scaled prior values.

#### Table S7. ABC observed summary statistics for modern samples

Table presenting the summary statistics used in ABC to compare models sampling only modern extant populations. *P*: population id number for correspondence on left matrices; $N_{ind}$: number of diploid individuals; *K*: mean number of alleles over loci; *H*: observed heterozygosity over loci; *GW*: Garza & Williamson (2001) index; *R*: Mean allelic range over loci; *prS*: Number of private polymorphic sites; *Pi*: Mean number of pairwise differences; $D_{\mu}^2$: square difference in mean STR allele length between pairs of populations, over loci; $F_{ST}$, $F_{IT}$, $F_{IS}$, *P*, sd: standard deviation. All statistics were estimated in arlsumstat.

| Modern samples only | | nDNA SSR | | | | mtDNA-dloop | | |
|---|---|---|---|---|---|---|---|---|
| Population | P | *K* | *H* | *GW* | *R* | *K* | *prS* | *Pi* |
| Western-Sahara | 1 | 2.89 | 0.35 | 0.41 | 10.2 | | 0 | 0.00 |
| Madeira | 2 | 1.53 | 0.12 | 0.27 | 13.0 | | 0 | 0.00 |
| Central-Eastern-Med | 3 | 4.63 | 0.42 | 0.30 | 16.8 | | 3 | 0.49 |
| Sd among pop | | 1.56 | 0.16 | 0.07 | 3.3 | | | 0.28 |
| Overall | | 5.37 | 0.53 | 0.31 | 19.7 | 5.0 | | |
| Overall sd | | 2.03 | 0.14 | 0.10 | 12.7 | | | |

| Population | nDNA $F_{ST}$ 1 | 2 | 3 | nDNA $D_{\mu}^2$ 1 | 2 | 3 |
|---|---|---|---|---|---|---|
| Western-Sahara | | 0.36 | 0.41 | | 5.2 | 11.3 |
| Madeira | 0.36 | | 0.49 | 5.2 | | 14.6 |
| Central-Eastern-Med | 0.41 | 0.49 | | 11.3 | 14.6 | |
| | $F_{IS}$ | $F_{IT}$ | $F_{ST}$ | | | |
| | 0.17 | 0.52 | 0.41 | | | |

| Population | mtDNA $F_{ST}$ 1 | 2 | 3 | mtDNA *Pi* 1 | 2 | 3 |
|---|---|---|---|---|---|---|
| Western-Sahara | | 0.00 | 0.76 | | 0.00 | 1.28 |
| Madeira | 0.00 | | 0.69 | 0.00 | | 1.28 |
| Central-Eastern-Med | 0.76 | 0.69 | | 1.28 | 1.28 | |
| | | | | *PI* | | |
| | | | | 0.49 | | |

#### Table S8. ABC observed summary statistics for modern and historic populations

Table presenting the summary statistics used in ABC to compare models including modern and historical populations. *P*: population id number for correspondence on left matrices; $N_{ind}$: number of diploid individuals; *K*: mean number of alleles over loci; *H*: observed heterozygosity over loci; *GW*: Garza & Williamson (2001) index; *R*: Mean allelic range over loci; *prS*: Number of private polymorphic sites; *Pi*: Mean number of pairwise differences; $D_{\mu}^2$: square difference in mean STR allele length between pairs of populations, over loci; FST, FIT, FIT, P, sd: standard deviation. All statistics were estimated in arlsumstat.

| 4 Modern pop + 1 Historical pop | | | nDNA SSR | | | | mtDNA dloop | | |
|---|---|---|---|---|---|---|---|---|---|
| Population | P | $N_{ind}$ | *K* | *H* | *GW* | *R* | *K* | *prS* | *Pi* |
| Western-Sahara | 1 | 93 | 2.89 | 0.35 | 0.41 | 10.2 | | 0 | 0.00 |
| Madeira | 2 | 10 | 1.53 | 0.12 | 0.27 | 13.0 | | 0 | 0.00 |
| Western-Med | 3 | 7 | 2.32 | 0.36 | 0.39 | 8.4 | | 2 | 1.73 |
| Central-Med | 4 | 26 | 3.21 | 0.44 | 0.32 | 12.5 | | 0 | 0.39 |
| Eastern-Med | 5 | 110 | 3.89 | 0.41 | 0.30 | 14.9 | | 1 | 0.40 |
| Sd among pop | | | 0.90 | 0.13 | 0.06 | 2.5 | | | 0.71 |
| Overall | | 246 | 5.53 | 0.53 | 0.31 | 19.9 | 6.0 | | |
| Overall sd | | | 2.09 | 0.14 | 0.11 | 13.0 | | | |

| Population | nDNA $F_{ST}$ 1 | 2 | 3 | 4 | 5 | nDNA $D_{\mu}^2$ 1 | 2 | 3 | 4 | 5 |
|---|---|---|---|---|---|---|---|---|---|---|
| Western-Sahara | | 0.36 | 0.24 | 0.42 | 0.42 | | 5.2 | 6.7 | 9.0 | 11.9 |
| Madeira | 0.36 | | 0.54 | 0.55 | 0.49 | 5.2 | | 10.3 | 10.7 | 15.5 |
| Western-Med | 0.24 | 0.54 | | 0.24 | 0.20 | 6.7 | 10.3 | | 10.1 | 8.3 |
| Central-Med | 0.42 | 0.55 | 0.24 | | 0.03 | 9.0 | 10.7 | 10.1 | | 2.7 |
| Eastern-Med | 0.42 | 0.49 | 0.20 | 0.03 | | 11.9 | 15.5 | 8.3 | 2.7 | |
| | $F_{IS}$ | $F_{IT}$ | $F_{ST}$ | | | | | | | |
| | 0.17 | 0.48 | 0.37 | | | | | | | |

| Population | mtDNA $F_{ST}$ 1 | 2 | 3 | 4 | 5 | mtDNA *Pi* 1 | 2 | 3 | 4 | 5 |
|---|---|---|---|---|---|---|---|---|---|---|
| Western-Sahara | | 0.00 | 0.68 | 0.97 | 0.80 | | 0.00 | 1.00 | 1.78 | 1.24 |
| Madeira | 0.00 | | 0.24 | 0.90 | 0.73 | 0.00 | | 1.00 | 1.78 | 1.24 |
| Western-Med | 0.68 | 0.24 | | 0.39 | 0.57 | 1.00 | 1.00 | | 1.59 | 1.57 |
| Central-Med | 0.97 | 0.90 | 0.39 | | 0.61 | 1.78 | 1.78 | 1.59 | | 1.02 |
| Eastern-Med | 0.80 | 0.73 | 0.57 | 0.61 | | 1.24 | 1.24 | 1.57 | 1.02 | |
| | | | | | | *PI* | | | | |
| | | | | | | 0.39 | | | | |

#### Table S9: ABC model fit

Table of statistics used to assess the model fit. **nr**: number of retained simulations, fit assessed using all summary statistics (**All stats**), reduced *N* components using a partial least square regression (**PLS**), using only subset of the statistics (**Selected stats**) or their PLS regression (**PLS on selected stats**).

#### Table S10: ABC structured model fit with selected statistics

Table of statistics used to assess the fit of structured models using summary statistics selected maximum entropy approach. **nr**: number of retained simulations.

#### Table S11: ABC parameter posterior estimates

Table of the parameter posteriors estimated in ABC. ABC-GLM posterior posteriors of the effective population size ($N_e$) and of the gene flow among populations (see stepping stone custom model in Figure S12) under the model M180 and M181, represented after ($N_0$: current population, $NM_0$ current gene flow), before ($N_1$: ancient / intermediate population , $NM_1$ ancient gene flow) the demographic event occurring at $T_1$ and before ($N_2$: ancient population, $NM_2$ ancient gene flow) the demographic event occurring at $T_2$ (M180 only). WS: Western-Sahara/Mauritania, MAD: Madeira, WM: Western-Mediterranean, CM: Central Mediterranean, EM: Eastern-Mediterranean. Natural scale time values have been multiplied by the generation time (10 years) to be expressed in years before present.

### Supporting Figures

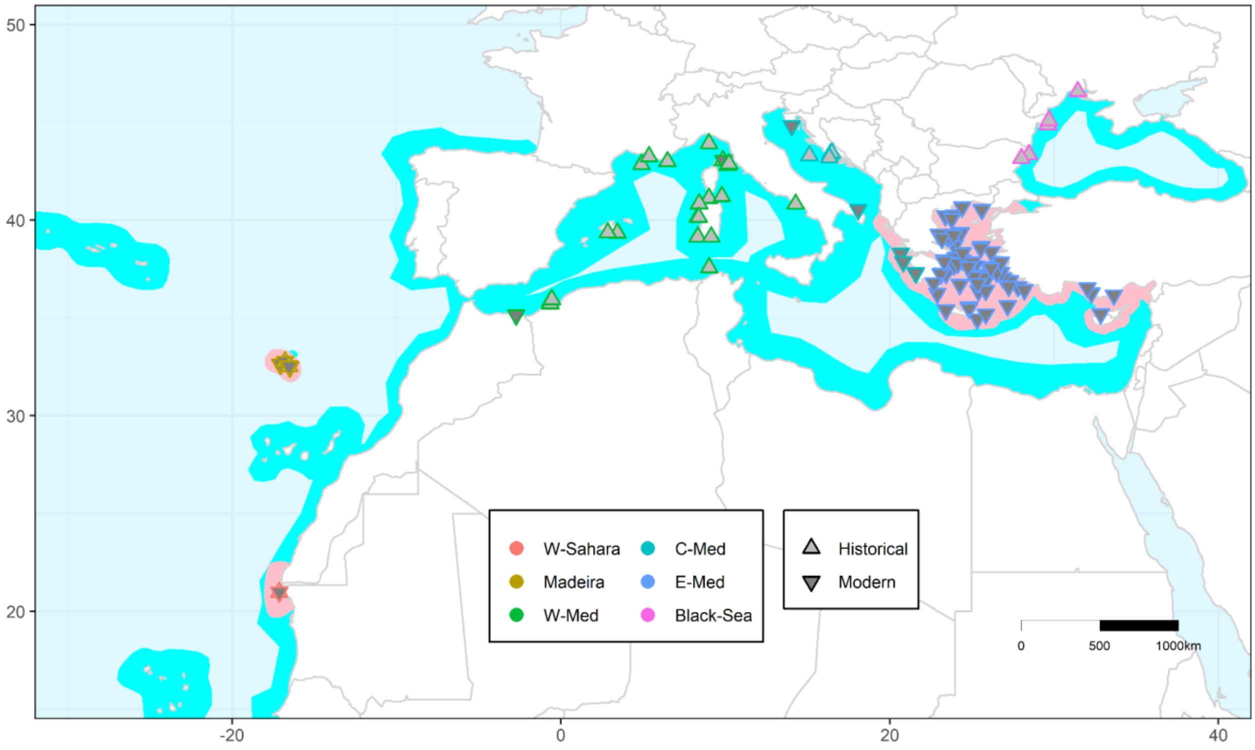


#### Figure S1. Map of the Mediterranean monk seals included in the study

The map presents all the samples (modern and historical) used in the study (sequenced at the CR1 mtDNA region and / or typed at the 19 microsatellites markers). The pink area represents *M. monachus*' current distribution, as defined by the IUCN in 2015 (Karamanlidis & Dendrinos, 2015). The cyan area represents the putative ancient distribution, according to (González, 2015). Detailed representations of samples historical *vs* modern and microsatellites *vs* mtDNA are presented in Figs S2. Numbers of samples in each locality and category (e.g. modern, historical) are given in Table 1. W-Sahara: Western-Sahara/Mauritania, W-Med: Western Mediterranean Sea, C-Med: Central Mediterranean Sea, E-Med: Eastern Mediterranean Sea.

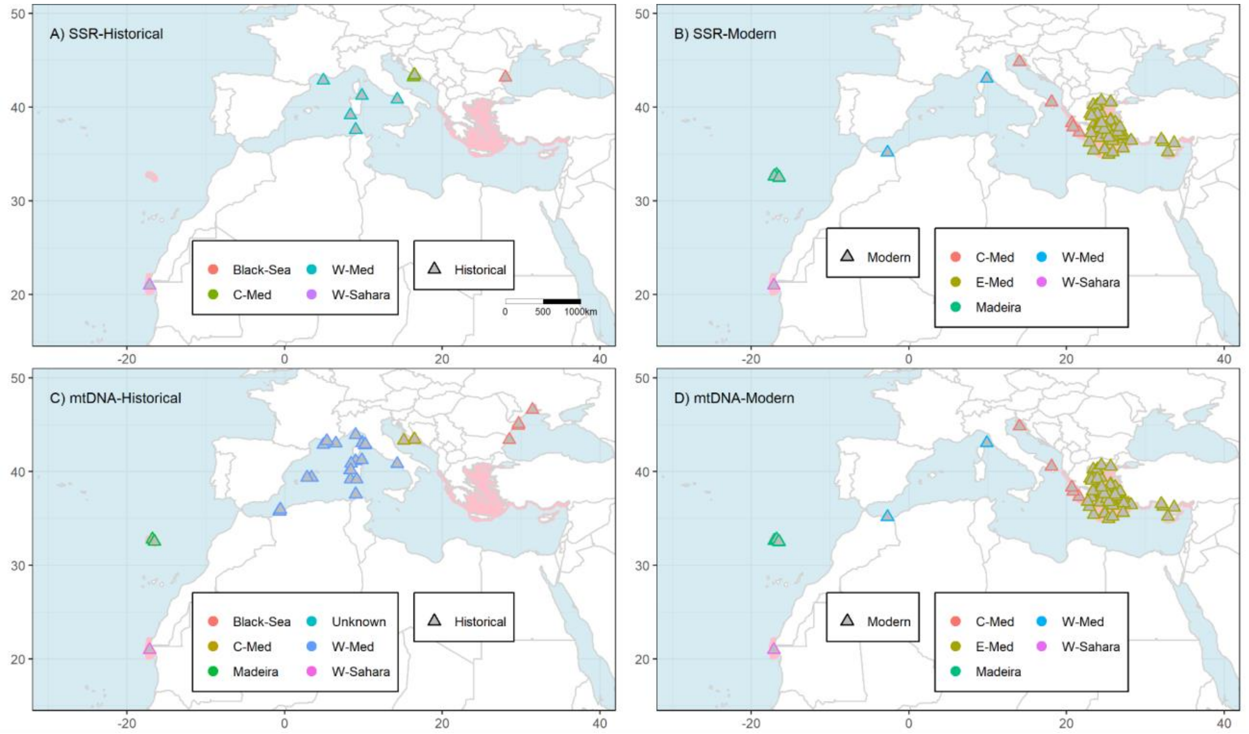


#### Figure S2. Detailed maps of the MMS samples included in the study

The figure depicts the Historical (A,C) and Modern (B,D) samples sorted by genotyping strategy (microsatellites A & B; and CR1 mtDNA: C & D). The pink area represents *M. monachus*' current distribution, as defined by the IUCN in 2015 (Karamanlidis & Dendrinos, 2015). Numbers of samples in each locality and category (e.g. modern, historical) are given in Tables S1a-b. SSR: microsatellites, W-Sahara: Western-Sahara/Mauritania, W-Med: Western Mediterranean Sea, C-Med: Central Mediterranean Sea, N-E-Med: Eastern Mediterranean Sea.

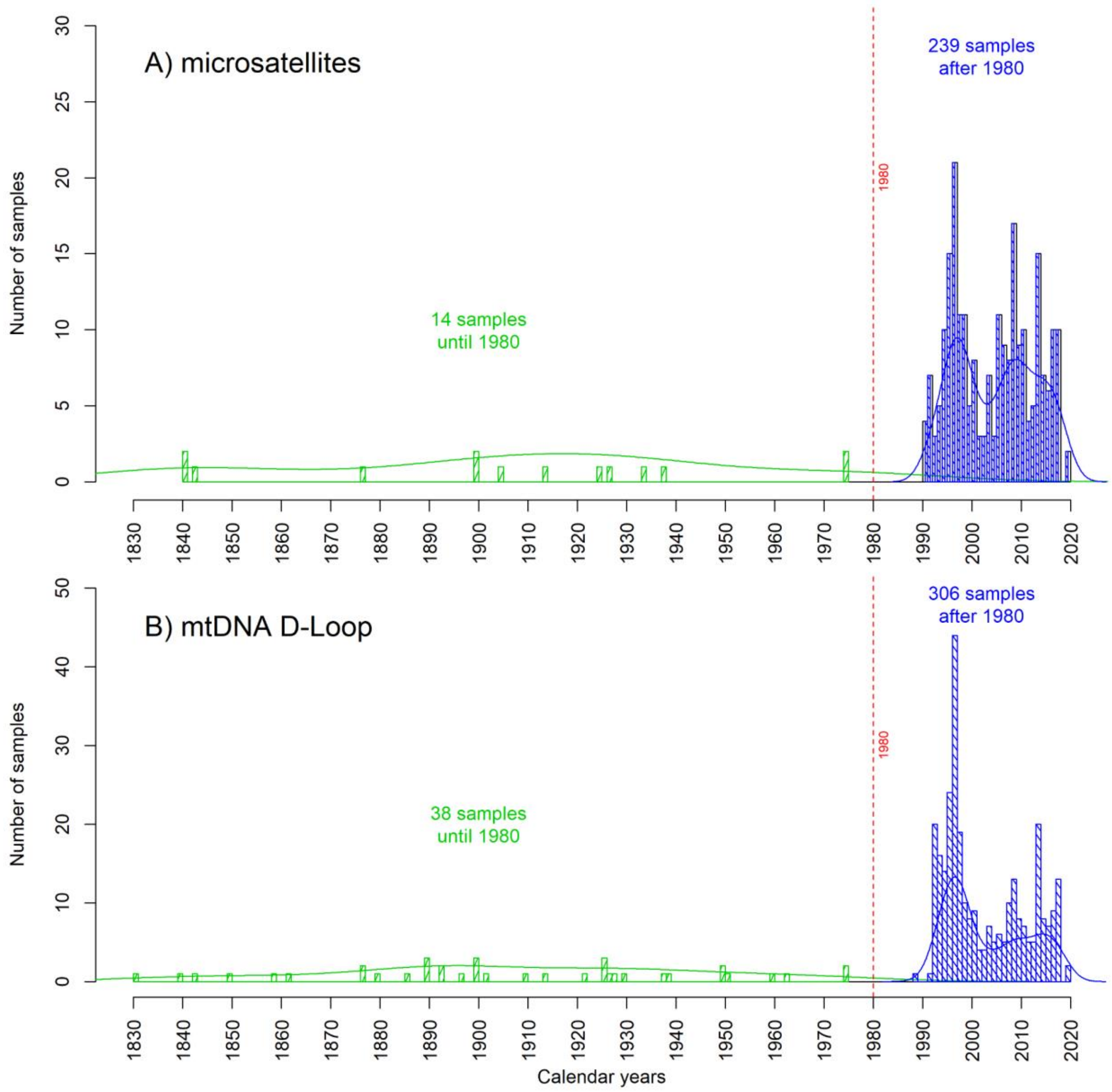


#### Figure S3. MMS sampling distribution through time

The figure shows the temporal distribution of genotyped samples included in the present study for microsatellites analyses (**A**) and mtDNA d-loop (**B**). The lines represent the estimated densities of the respective sampling.

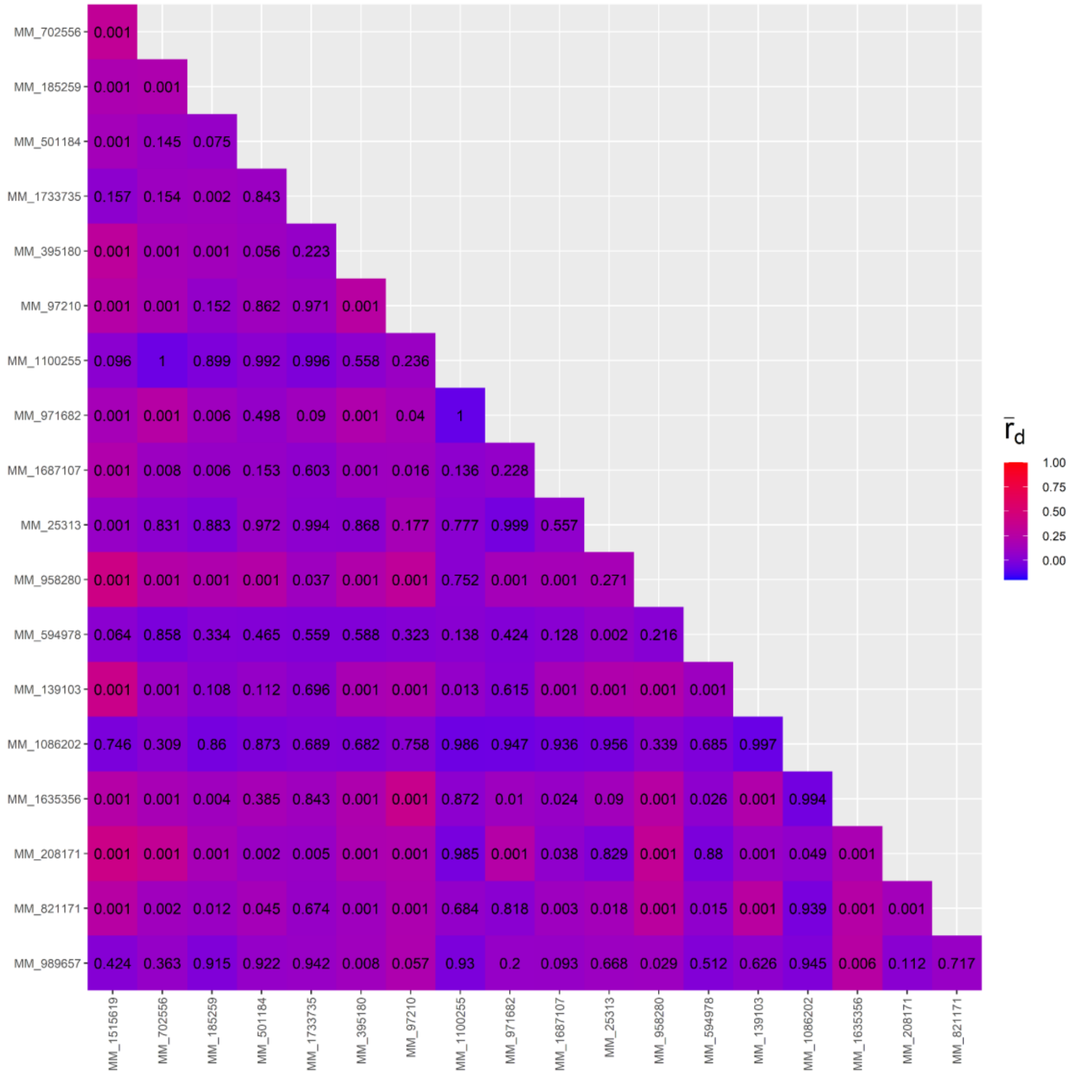


#### Figure S4. Microsatellites linkage disequilibrium across all samples

The Agapov and Burt $r_d$ values of linkage disequilibrium among microsatellites loci (estimated across all 253 individuals included in the study) are represented with the red to blue color scale, their associated *p*-values are noted in their respective cells, x and y axis labels represent the microsatellites locus ID. High (and significant) $r_d$ value (cf. color scale) indicates a possible linkage between the makers. Despite the low diversity of the markers and the MMS structure pattern (Fig. 1) that both confound the LD estimates none of the loci pairs shows high LD values. Furthermore none of the LD values stay significant when estimated at the regional scale (Fig. S5-6).

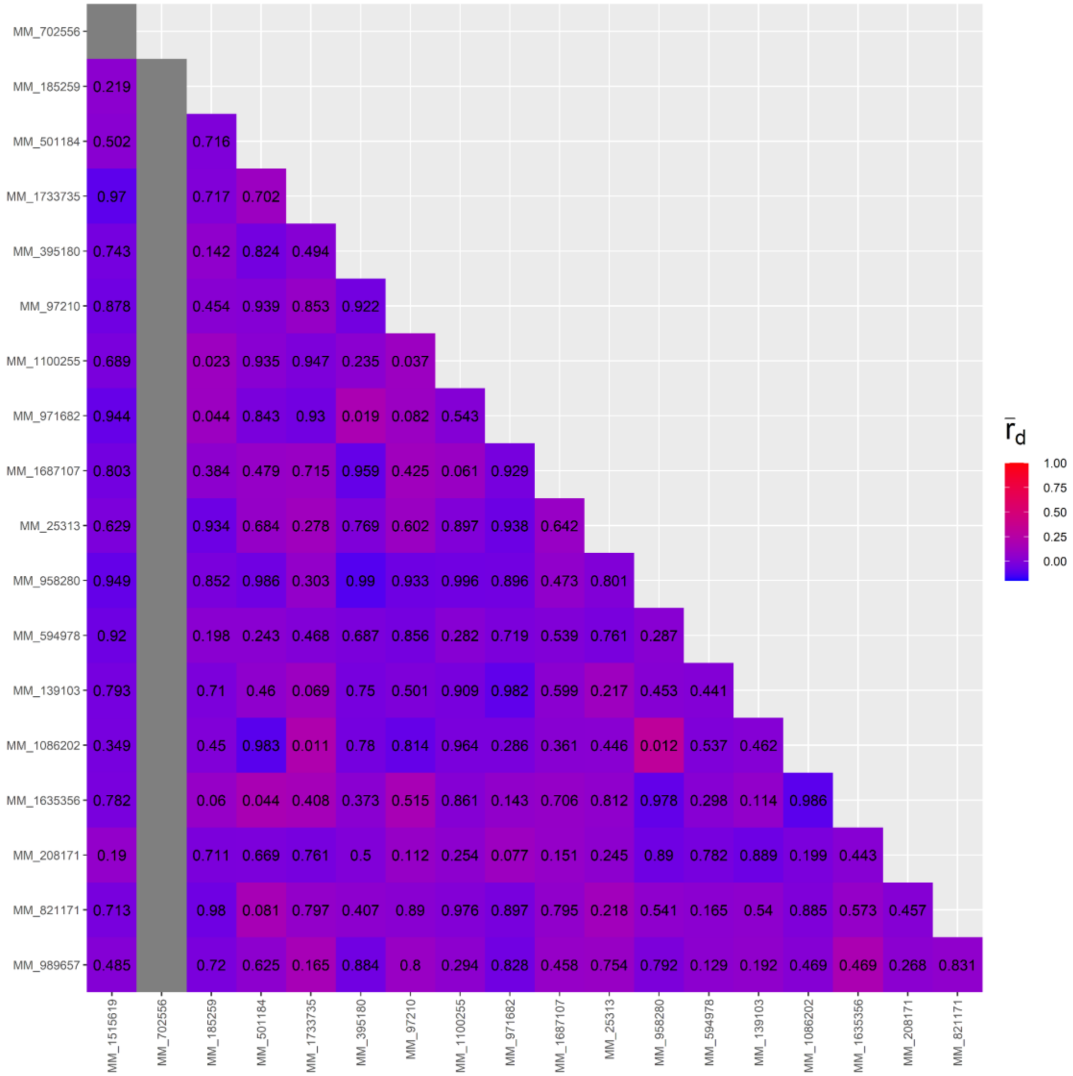

#### Figure S5. Microsatellites linkage disequilibrium WS samples

The Agapov and Burt $r_d$ values of linkage disequilibrium among microsatellites loci (estimated across the 96 Western-Saharan individuals) are represented with the red to blue color scale, their associated *p*-values are noted in their respective cells, x and y axis labels represent the microsatellites locus ID. High (and significant) $r_d$ value (cf. color scale) indicates a possible linkage between the makers. At the level of the western-Sahara/Mauritiana population, and despite the low diversity of the markers that confound the LD estimates, none of the loci pairs shows high LD values. The few pairs comparisons that show significant trends, are however not confirmed with analyses at the global level or at the eastern-Mediteranean scale.

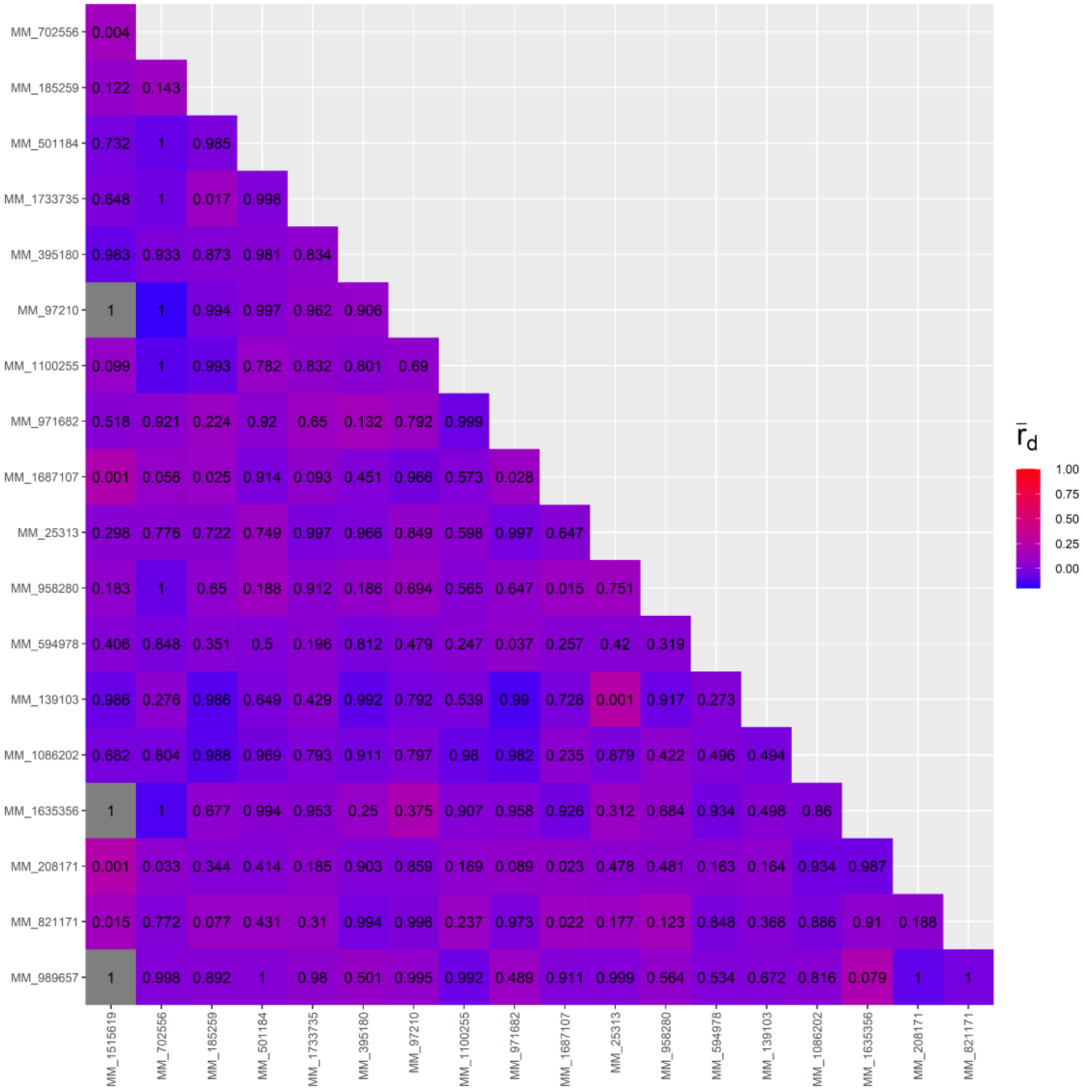


### Figure S6. Microsatellites linkage disequilibrium EM samples

The Agapov and Burt $r_d$ values of linkage disequilibrium among microsatellites loci (estimated across the 148 eastern-Mediteranean individuals) are represented with the red to blue color scale, their associated *p*-values are noted in their respective cells, x and y axis labels represent the microsatellites locus ID. High (and significant) $r_d$ value (cf. color scale) indicates a possible linkage between the makers. At the level of the western-Sahara/Mauritania population, and despite the low diversity of the markers that confound the LD estimates, none of the loci pairs shows high LD values. The few pairs comparisons that show significant trends, are however not confirmed with analyses at the global level or at the Western-Sahara/mauritanian scale.

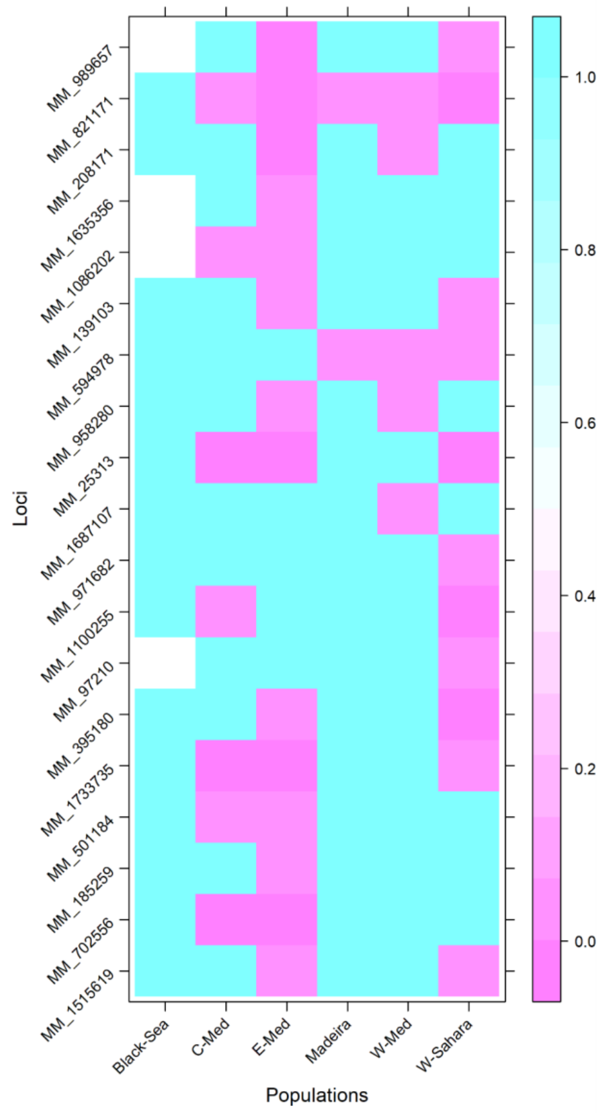


#### Figure S7. Microsatellites Hardy-Weinberg equilibrium

Representation of the Hardy-Weinberg equilibrium test *p*-values estimated per population using 1 000 Monte Carlo permutations of alleles (Guo & Thompson, 1992). Note, all non-significant *p*-values (> 0.05) are shown in light blue.

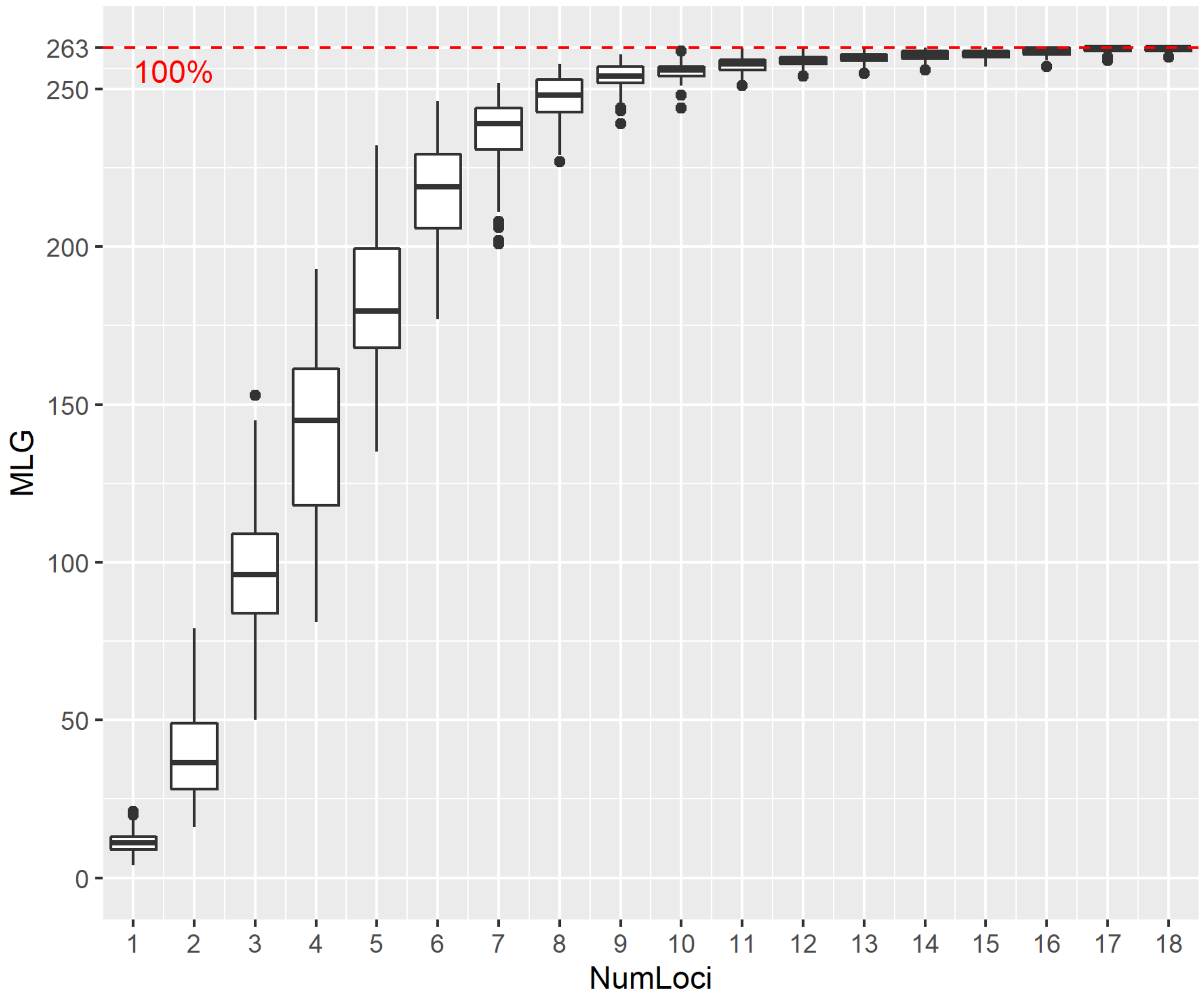


#### Figure S8. Microsatellites genotype accumulation curve

Genotype accumulation curve used for determining the minimum number of loci necessary to discriminate between individuals in a population. This function randomly samples loci without replacement and counts the number of multilocus genotypes observed. Here at ~>10 loci we start to fairly discriminate individuals from each other. In other words, we should filter-out genotypes with missing data at nine or more loci. Numloci: Number of loci; MLG: Multi locus genotype.

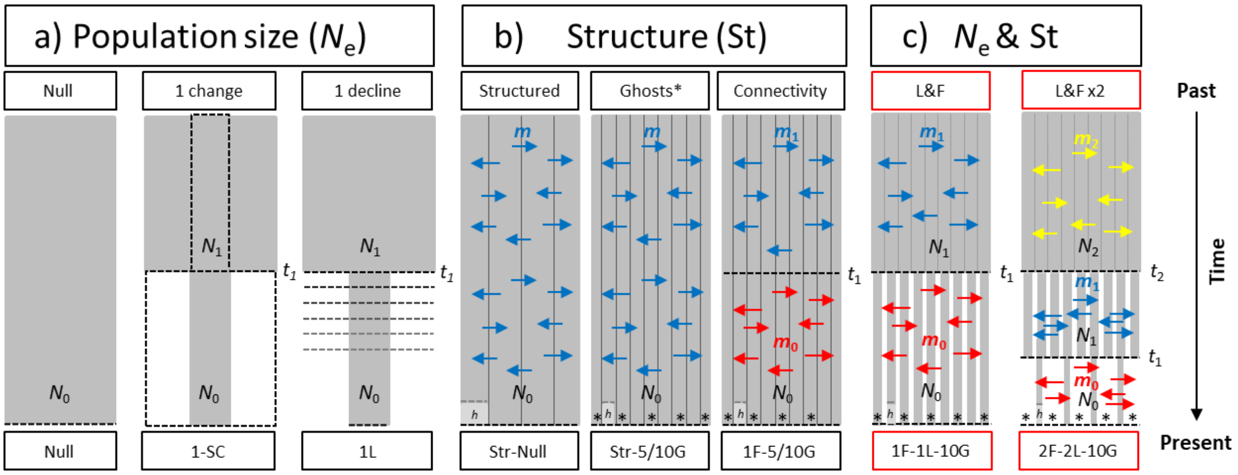


#### Figure S9: Summary of demographic models tested using ABC.

Main demographic scenarios compared within the ABC framework. Schematic representation of the tested models. Models in (**a**) with panmictic population were built to test the effect of habitat loss alone, without fragmentation. These scenarios range from the null model with no population size change, to the one loss model (1L) that models one population size decline. The scenario 1-SC allows for one population size change (growth or decline) see Fig. S10 for further details. In (**b**), we modeled stable-size structured populations with constant migration rates among populations (Str-Null). These additionally included (i) 5 or 10 unsampled ghost populations (Str-5/10G) mimicking the extinct populations of (Black Sea, Canarias, Azores, etc), and (ii) a change in connectivity at $t_1$ (1F-5G), mimicking the loss of gene flow among major colonies, see Figs S11-12 for further details. The last scenarios (**c**) combine population size change, structure, ghost populations, and changes in connectivity, to model populations that suffered one (1F-1L-5G) or several (e.g. 2F-2L-5G) events of decline and fragmentation, see Fig. S13. The most supported models are highlighted with a red box. In **b** and **c** the historic Western-Mediterranean population sampled ~140 years in the past is represented by an '*h*'. Subpopulation numbers and positions, as well as number and direction of arrows of gene flow between demes are an arbitrary representation and do not represent the exact number of populations or of gene flow events, for parameterization values see Table S5. SC: size change, L: loss, Str: structure, G: ghost, F: fragmentation, L&F: loss and fragmentation, $N_n$: population size, $t_n$: time of event, $m_n$: migration rate.

Supporting information for: The antique genetic plight of the Mediterranean monk seal (*Monachus monachus*). J. Salmona et al., in revision. doi:10.1098/rspb.2022-0846.

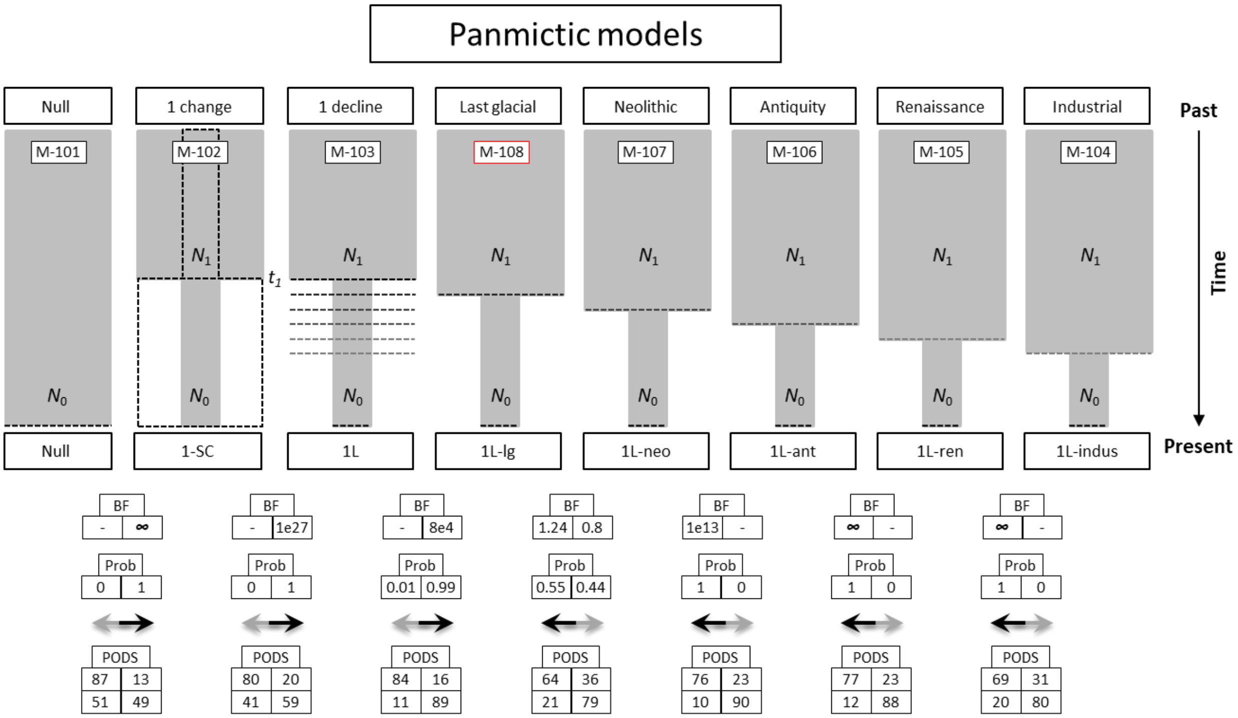


#### Figure S10: ABC panmictic model choice

Schematic representation of the ABC-GLM model testing procedure, conducted across panmictic models to assess the effect of population fluctuations alone. The figure illustrates that models with one population decline (1L) are more supported than models including both decline or growth (1-SC), or stationary models (NULL). Furthermore, the scenario allowing for the largest population decline time window (1L) is more supported than any other models narrowing this window to specific time hypotheses (“Last glacial”, “Neolithic”, “Antiquity”, “Renaissance” and “Industrial”. Top box: Bayes factor (BF), middle box: model posterior probabilities (Prob), for each pair of models. The bottom box illustrates the power to distinguish between the two compared models as evaluated in a cross-validation procedure with 1,000 validations for each model (PODS), with the upper left and lower right boxes showing the correct model assignments for left and right models respectively. $N_0$: current effective population size, $N_1$: ancient effective population size, $t_1$: time of a population size change. The selected model is highlighted with a red rectangle.

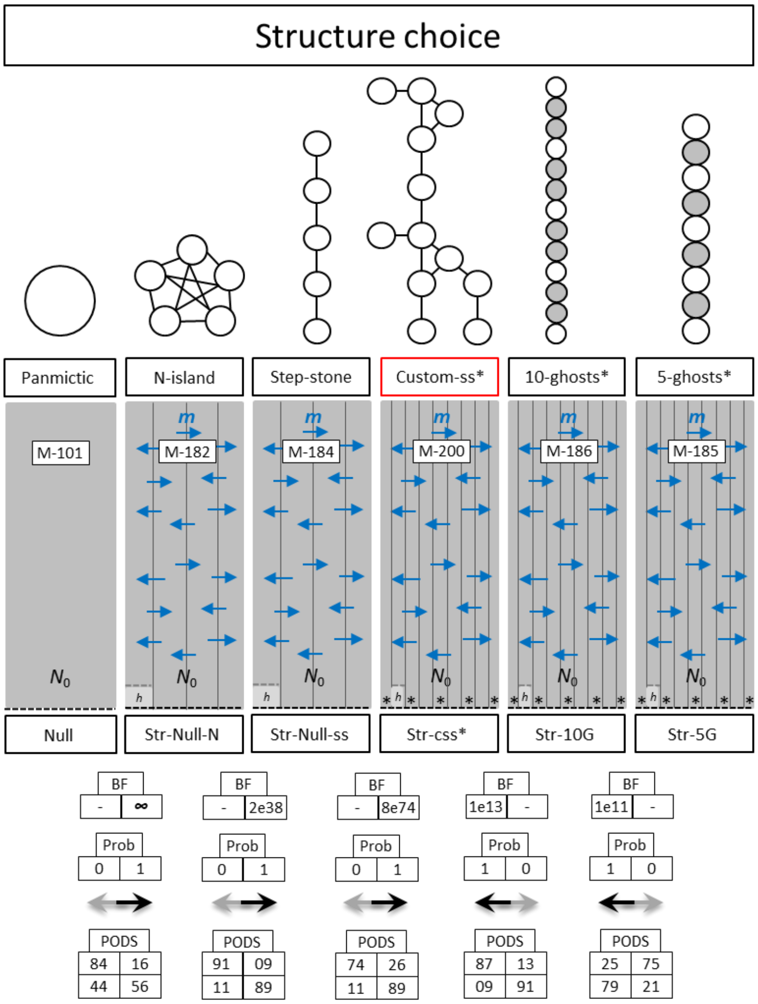


#### Figure S11: ABC structure choice

Schematic representation of the ABC-GLM model testing procedure, conducted across distinct population structure types. The structure types tested range from a model with no population structure (panmixia), to models with structure and ghost unsampled populations. With strong support, the custom stepping-stone model developed to approximate the possible structure of extant and extinct MMS populations, shows a better fit to the data when compared to panmictic, *n*-island, and linear stepping stone models. Top box: Bayes factor (BF), middle box: model posterior probabilities (Prob), for each pair of models. The bottom box illustrates the power to distinguish between the two compared models as evaluated in a cross-validation procedure with 1,000 validations for each model (PODS), with the upper left and lower right boxes showing the correct model assignments for left and right models respectively. $N_0$: current effective population size, *m*: migration rate between demes, *: ghost unsampled population. The best fitting and selected models are highlighted with darker arrows and red rectangles respectively.

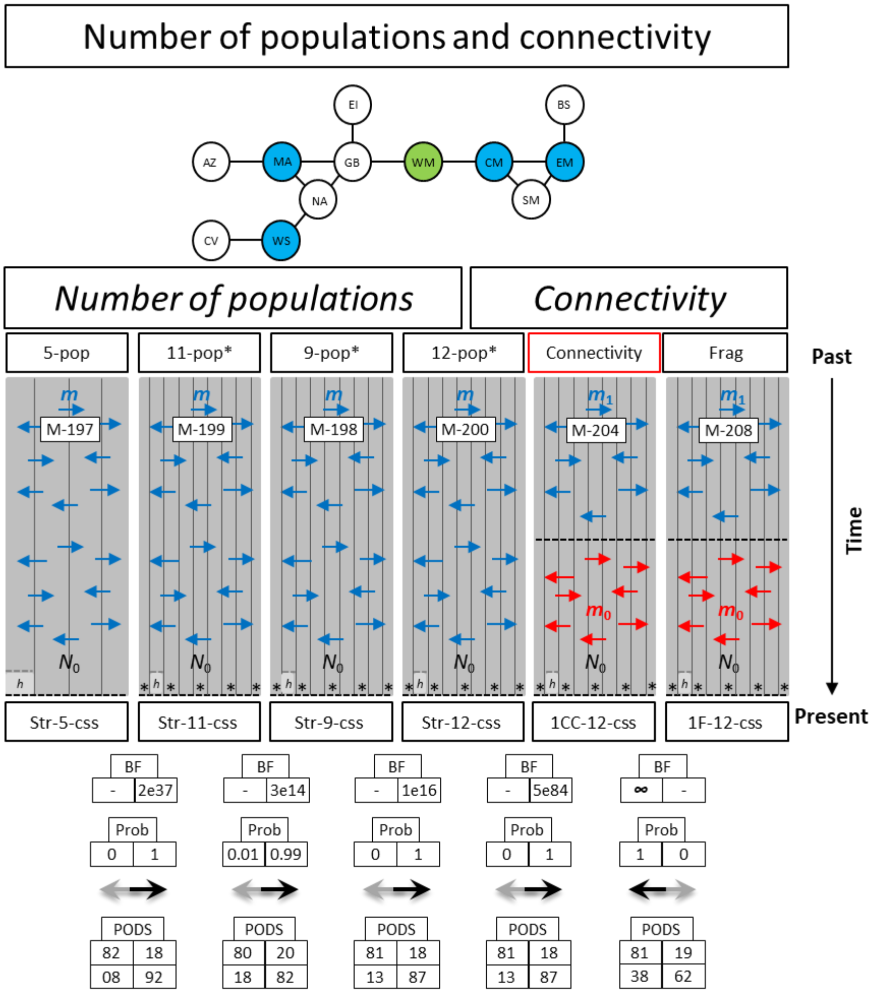


#### Figure S12: ABC structured stationary model choice

Upper plot: schematic representation of the custom stepping-stone model developed to approximate the possible structure of extant and extinct MMS populations. Lower plot: schematic representation of the ABC-GLM model choice procedure, conducted across structured scenarios, to assess the effect of the number of populations (5-9-11-12) and the type of connectivity changes (large prior [Connectivity] or decrease [Frag]). Top box: Bayes factor (BF), middle box: model posterior probability (Prob). The bottom box illustrates the power to distinguish between the two compared models as evaluated in a cross-validation procedure with 1,000 validations for each model (PODS), with the upper left and lower right boxes showing the correct model assignments for left and right models respectively. $N_0$: current effective population size, *m*: migration rate between demes, *: ghost unsampled population. The best fitting and selected models are highlighted with darker arrows and red rectangles, respectively. 5-pop: 5 sampled pops, 12-pop: all populations represented in the upper cartoon, 9-pop: EI, AZ and CV not included, 11-pop: EI not included. AZ: Azores, MA: Madeira, CV: Cape Verde, WS: Cabo Blanco, NA: North-East Africa, EI: Eastern Iberia, GB: Gibraltar, WM: Western Mediterranean Sea, CM: Central Mediterranean Sea, SM: South-Eastern Mediterranean Sea, EM: Eastern Mediterranean Sea, BS: Black Sea.

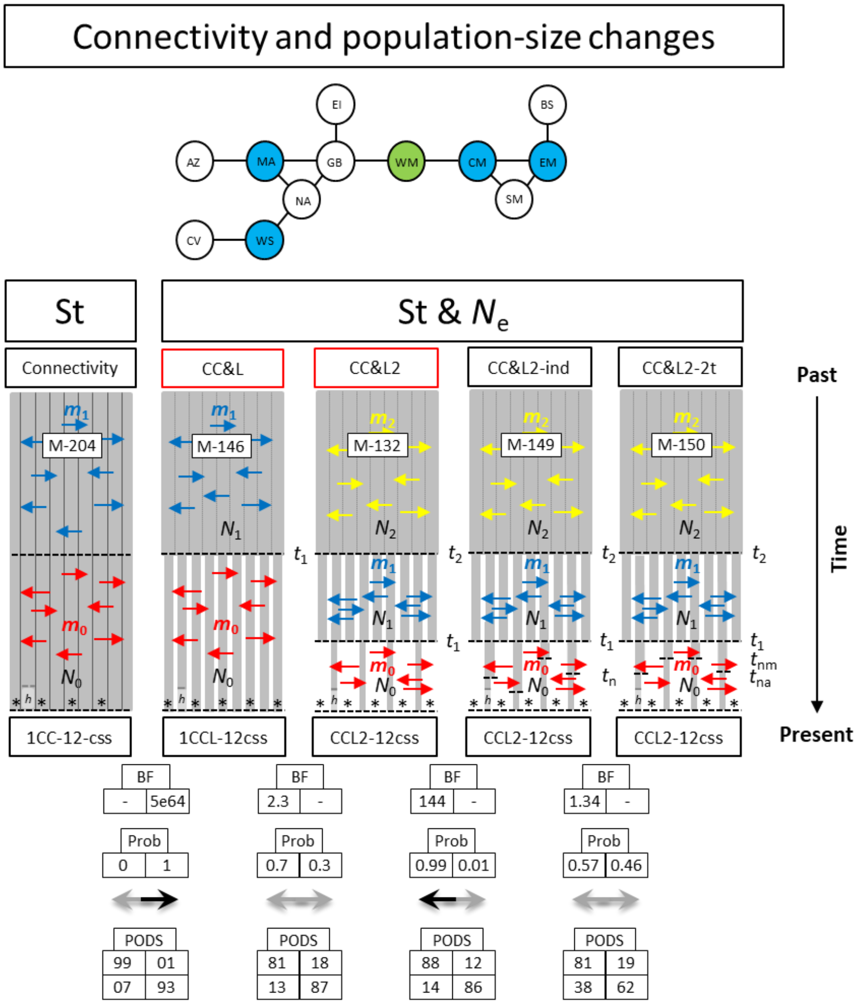


### Figure S13: ABC complex model choice

Upper plot: schematic representation of the custom stepping-stone model developed to approximate the possible structure of extant and extinct MMS populations. Lower plot: schematic representation of the ABC-GLM model choice procedure, conducted across structured scenarios including population size decrease(s), to assess the effect of the number of populations decrease (1-3) and of connectivity changes. Top box: Bayes factor (BF), middle box: model posterior probability (Prob). The bottom box illustrates the power to distinguish between the two compared models as evaluated in a cross-validation procedure with 1,000 validations for each model (PODS), with the upper left and lower right boxes showing the correct model assignments for left and right models respectively. $N_0$: current effective population size, *m*: migration rate between demes, *: ghost unsampled population. The best fitting and selected models are highlighted with darker arrows and red rectangles, respectively. AZ: Azores, MA: Madeira, CV: Cape Verde, WS: Cabo Blanco, NA: North-East Africa, EI: Eastern Iberia, GB: Gibraltar, WM: Western Mediterranean Sea, CM: Central Mediterranean Sea, SM: South-Eastern Mediterranean Sea, EM: Eastern Mediterranean Sea, BS: Black Sea.

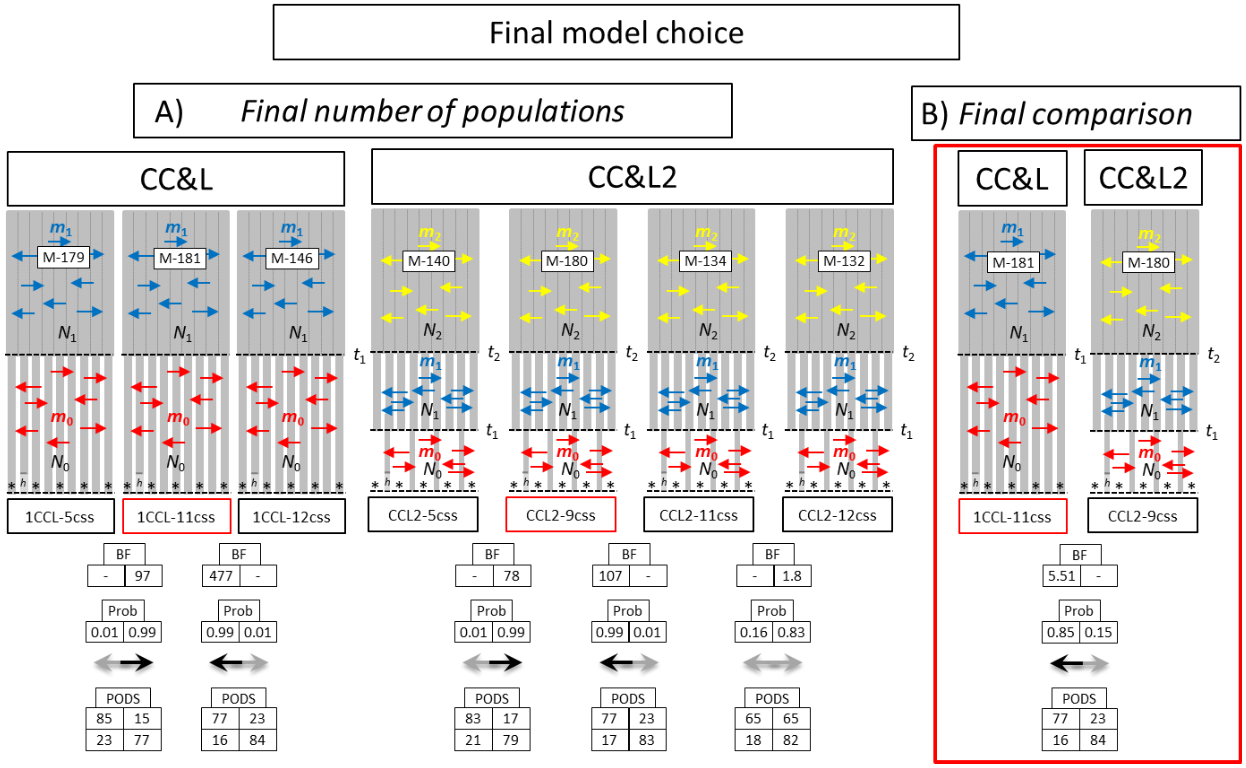


### Figure S14: ABC final model choice

Schematic representation of the ABC-GLM model choice procedure, conducted across structured scenarios including population size decrease(s), to assess the effect of the number of populations decrease (1-2), of populations (9-12) and of connectivity changes (1-2). Top box: Bayes factor (BF), middle box: model posterior probability (Prob). The bottom box illustrates the power to distinguish between the two compared models as evaluated in a cross-validation procedure with 1,000 validations for each model (PODS), with the upper left and lower right boxes showing the correct model assignments for left and right models respectively. $N_0$: current effective population size, $m$: migration rate between demes, *: ghost unsampled population. The best fitting and selected models are highlighted with darker arrows and red rectangles respectively.

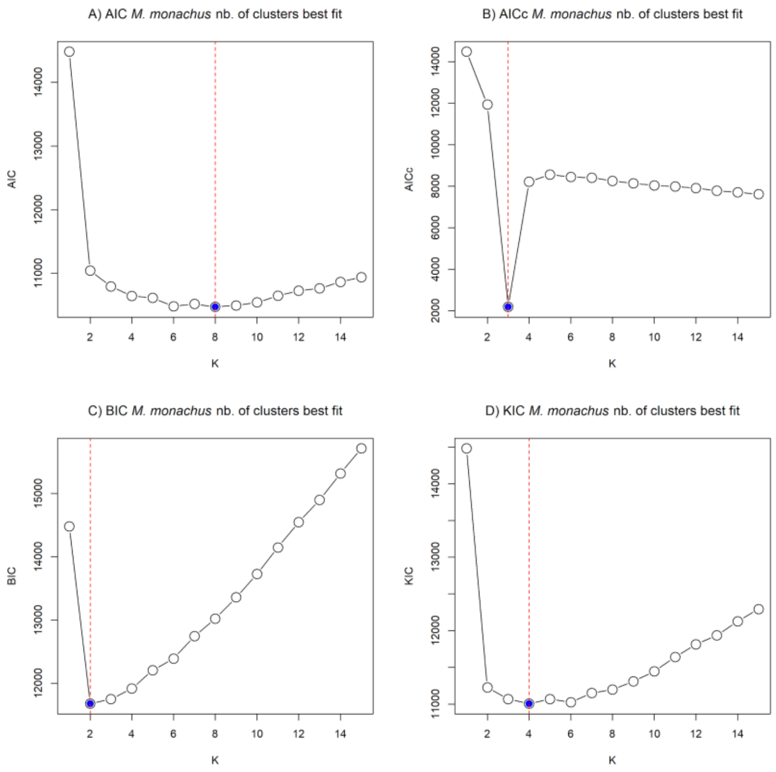


### Figure S15: Most likely number of panmictic clusters in MMS

The four goodness of fit estimates (A) AIC, (B) AICc, (C) BIC and (D) KIC used to assess the most likely number of clusters (K) in *M. monachus* show varying responses for numbers between 2 and 8 and with a narrower window of values for AICc, BIC and KIC between 2 and 4. Among the four goodness of fit metrics used to assess the most likely number of clusters (K) in M. monachus, three point to values (K=2 [BIC], K=3 [AICc] and K=4 [KIC], Fig. S14) exhibiting geographically coherent results (Fig. S15), the last one pointing to little interpretable results (K=8 [AIC]). The lack of congruence among these metrics is expected in systems exhibiting continuous genetic diversity (e.g. Isolation by distance), such as *M. monachus*.

Supporting information for: The antique genetic plight of the Mediterranean monk seal (*Monachus monachus*). J. Salmona et al., in revision. doi:10.1098/rspb.2022-0846.

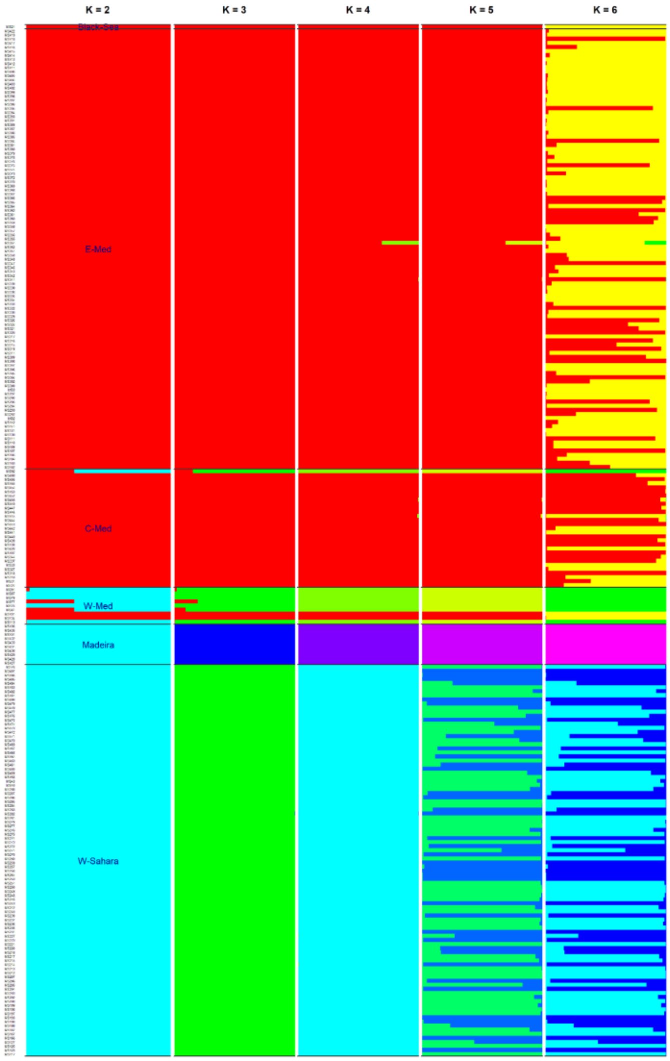


#### Figure S16: MMS genetic clustering

The individual membership coefficient estimated using snapclust (Tonkin-Hill *et al.*, 2019) represented by horizontal bar plots, showing that applying little supported numbers of clusters superior to four ($K \geq 5$, Fig. S14), divides the W-Saharan ($K = 5$) or E-Mediterranean samples ($K = 6$) into little-coherent sub-populations. K: number of investigated panmictic clusters.

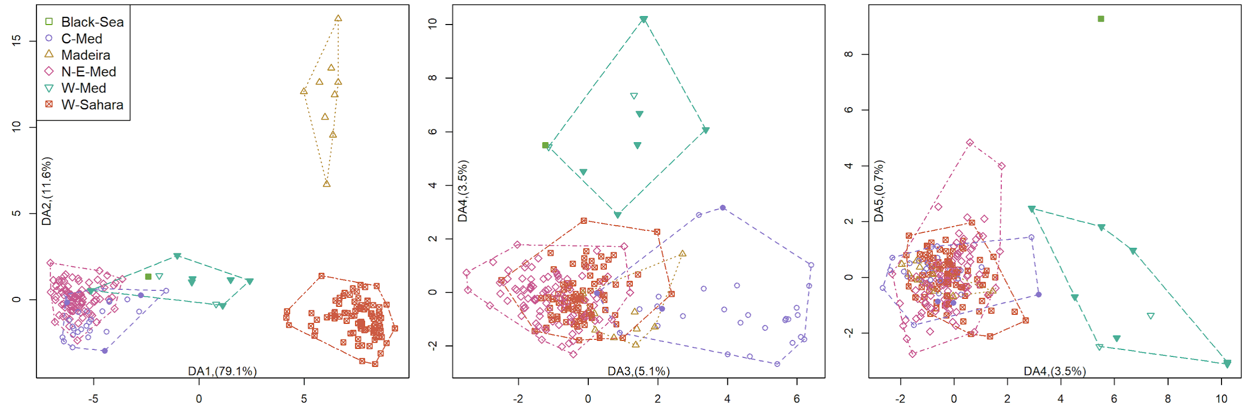


#### Figure S17: DAPC representation of microsatellites structure

The figure presents *M. monachus*' genetic structure revealed by microsatellite markers. The sample representation on the five axes of discriminant analysis on principal components DAPC summarize the genetic diversity of monk seals. Full and empty symbols represent historical and modern samples respectively. The figure shows (i) a strong West-East cline of differentiation on the first axis (left panel); (ii) a clear differentiation of Madeira samples on the second axis (left panel) suggesting a substantial differentiation of the Madeira island population with regards to the coastal populations.

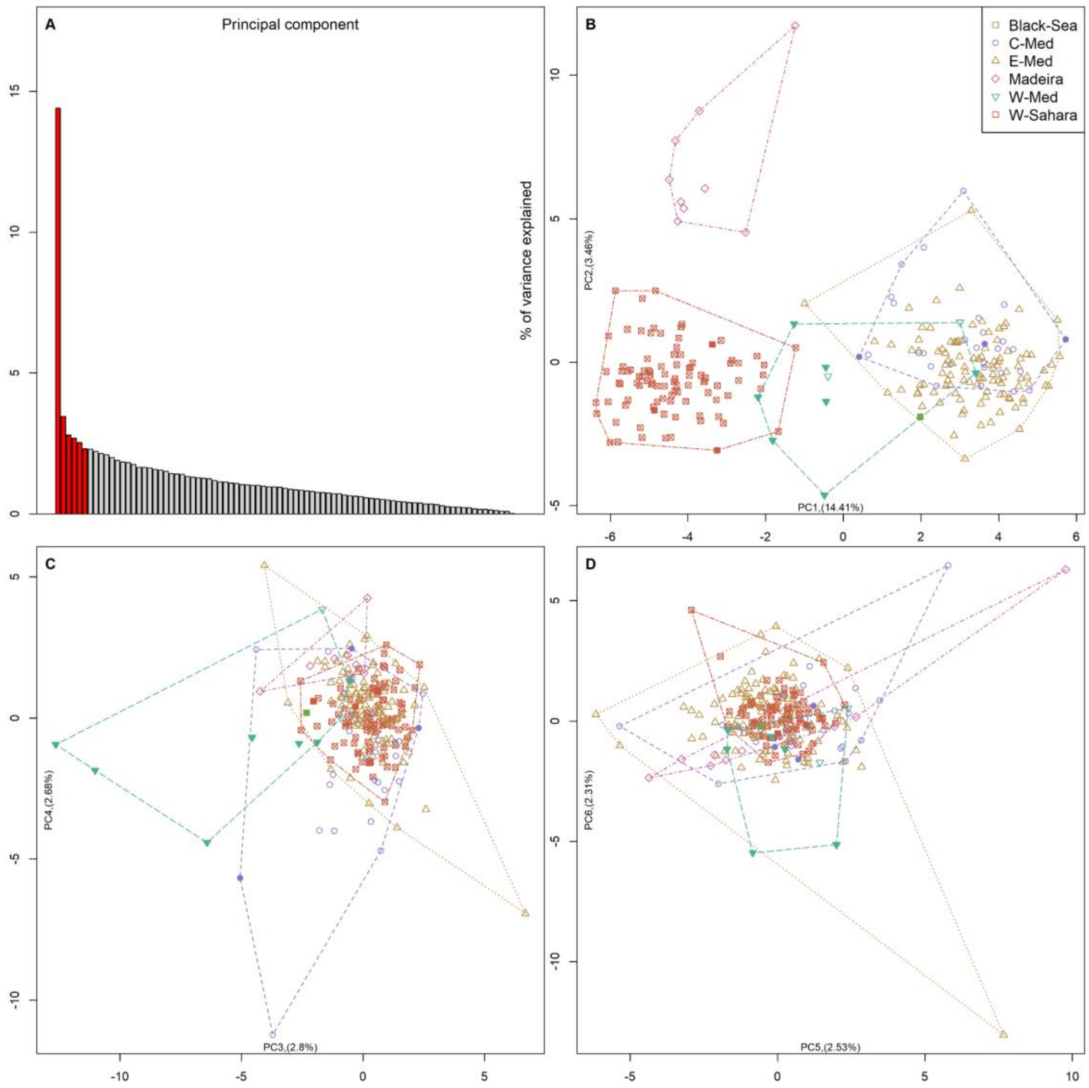


### Figure S18: PCA representation of microsatellites structure.

The figure presents *M. monachus*' genetic structure revealed by a principal component analysis of the microsatellite markers. The relative contribution of each component represented in (**A**), shows that the first two axes have a consequent contribution. The six first represented axes are highlighted in red. The sample representation on the first two axes of principal components analysis (**B**) summarizes the genetic diversity of monk seals and echoes particularly well the geography of the sampled regions. It shows a strong West-East cline of differentiation on the first axis and a clear differentiation of Madeira and Western-Sahara/Mauritania on the second axis (**B**). The interpretation of bidimensional representation of components 3-6 explaining lower amounts of variance (**C-D**) is not trivial. In **B-D**, full and empty symbols represent historical and modern samples respectively. W-Sahara: Western-Sahara/Mauritania, W-Med: Western Mediterranean Sea, C-Med: Central Mediterranean Sea, E-Med: Eastern Mediterranean Sea.

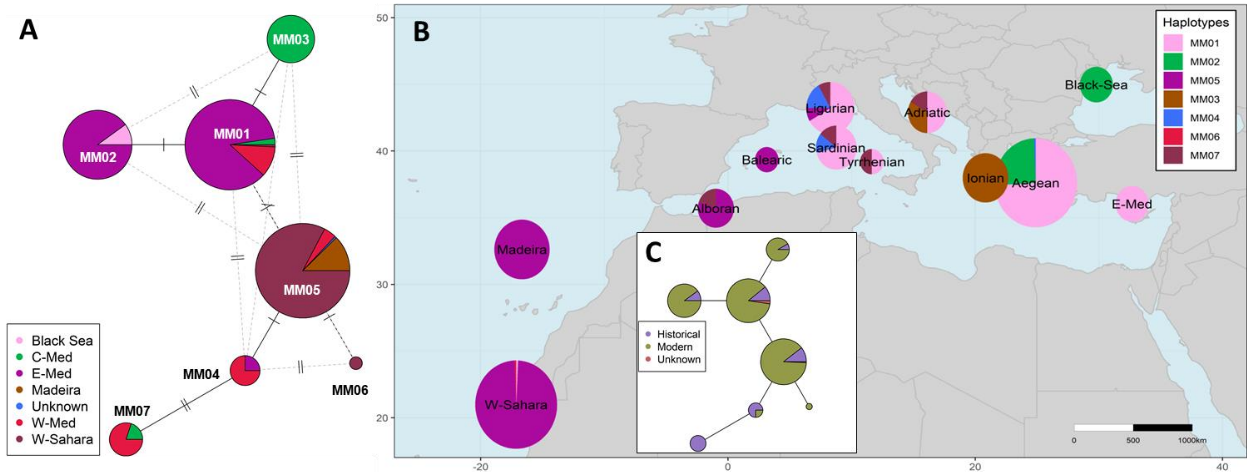


### Figure S19. *Monachus monachus* mitochondrial genetic diversity.

The figure presents *M. monachus*' D-loop genetic structure. The haplotype network (**A**-**C**) and their geographic representation (**B**) illustrates the low mitochondrial genetic diversity of Mediterranean monk seals (**A**), its changes over time (**C**), as well as the little overlap between the distribution of eastern and western haplotypes (**A**-**B**). In **A** and **C**, the haplotype pie charts show their relative representation across the main geographic areas (in **A**), and across time (in **C**), the lines show the main and secondary (dotted) relationship, and the bars the number of mutations, between haplotypes. In **B**, the population pie charts show their relative composition in d-loop haplotypes. In all three plots the haplotypes names follow Karamalidis et al. 2016 and Gaubert et al., 2019, and their size is proportional to the log of the number of individuals included (times $3.5e^{-2}$ in A and C, and $5.6e^{-1}$ in B). Complementary illustrations of the evolution of haplotype diversity over time are presented in the figures S19-20. W-Sahara: Western-Sahara/Mauritania, W-Med: Western Mediterranean Sea, C-Med: Central Mediterranean Sea, E-Med: Eastern Mediterranean Sea, Unknown: sample with unclear origin.

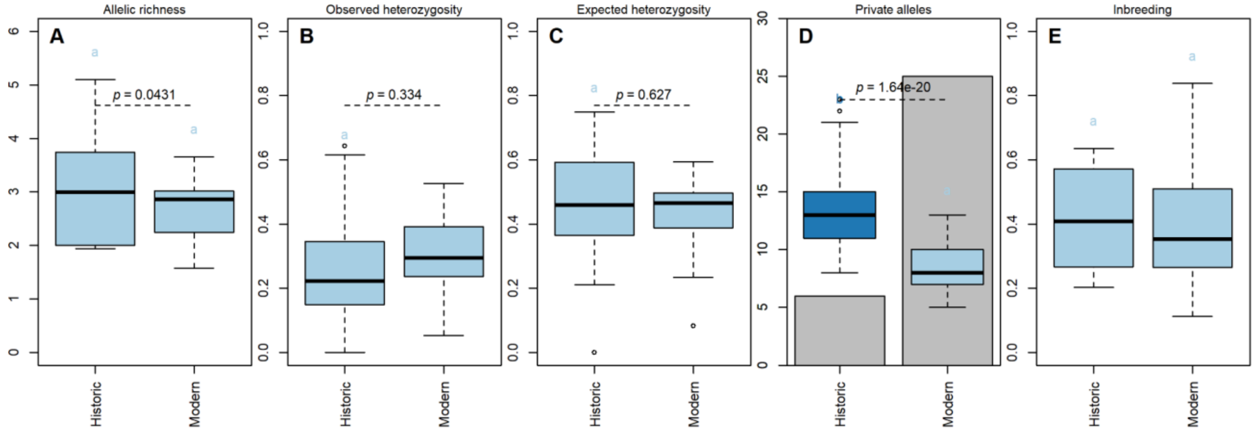


#### Figure S20. Temporal evolution of MMS genetic diversity.

Boxplot representation of the microsatellites (**A**) allelic richness ($A_R$), (**B**) observed ($H_O$), and (**C**) expected ($H_E$) heterozygosity, (**D**) boxplot of mean private alleles ($P_A$) over 500 resampling of the smallest sample size, and barplot of the counted number of private alleles ($P_A$), and (**E**) boxplot of the individual inbreeding ($F$) estimates over time, expressed for modern (>1979) and historical (<1980) samples respectively. The letters and boxplot-colors illustrate the Tukey post-hoc group assignment per period. Horizontal dashed-bars in **A-C** illustrate the *p-value* of the Student's t-test comparing the distribution of markers estimates among periods. To overcome the sampling bias between time periods, $H_O$ and $H_E$ and $P_A$ estimates (**B**-**D**) were averaged across 500 random subsamples of size 14 (size of the smallest group).

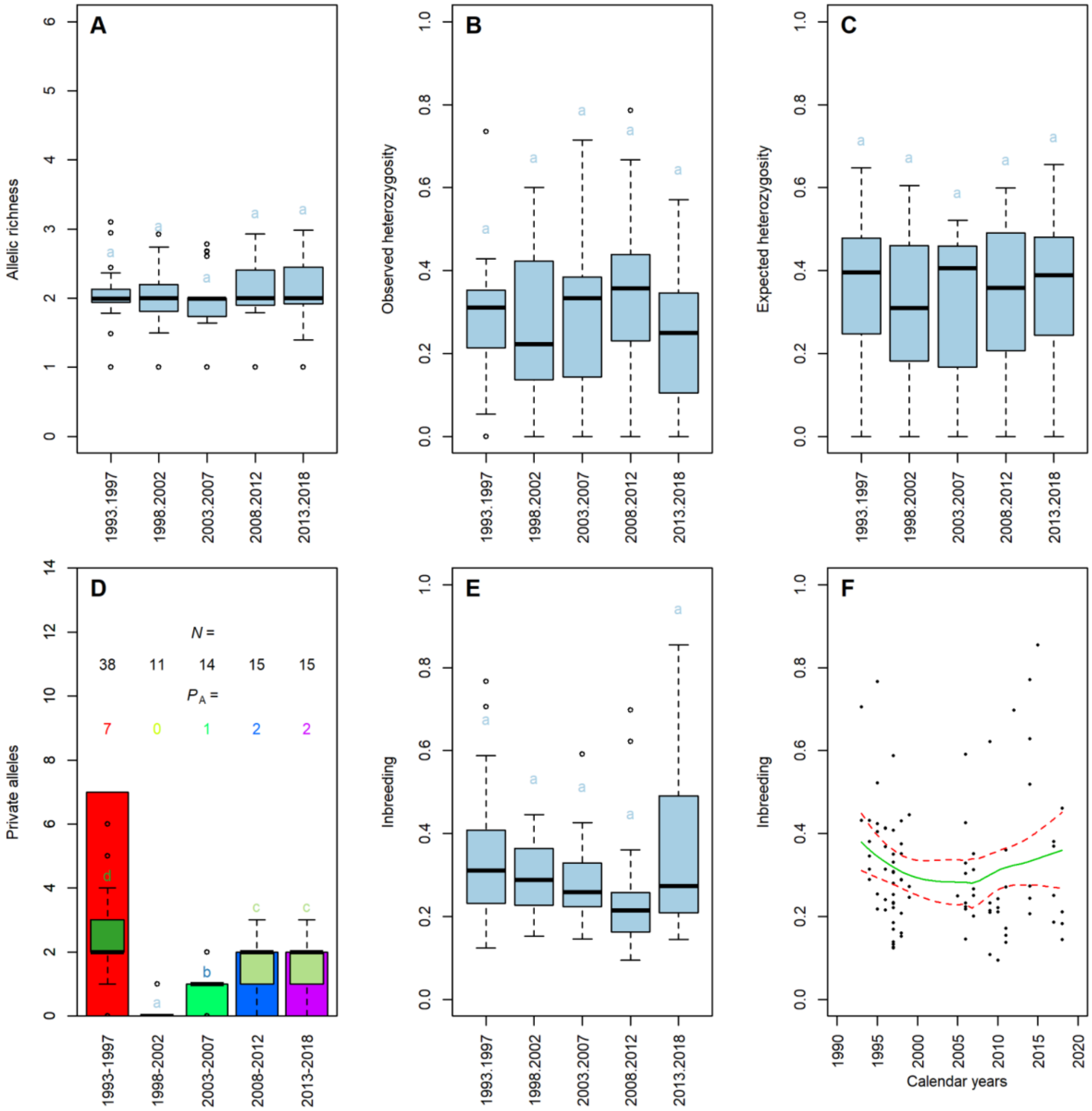


#### Figure S21: Temporal evolution of the Western-Saharan/Mauritania genetic diversity

Boxplot representation of the microsatellites (**A**) allelic richness ($A_R$), (**B**) observed ($H_O$), and (**C**) expected ($H_E$) heterozygosity, (**D**) barplot of the number of private alleles ($P_A$), and boxplot (**E**) and Smooth Local Polynomial Regression Fitting (**F**) representations of the individual inbreeding (*F*) estimates over time. In **A-E** the data is arbitrarily splitted in 5 years time intervals. The letters and boxplot-colors in **A-C** and **E** illustrate the Tukey post-hoc group assignment per period. In **F** the green continuous and red dotted lines represent the Smooth Local Polynomial Regression Fitting and its 95% confidence-interval, respectively.

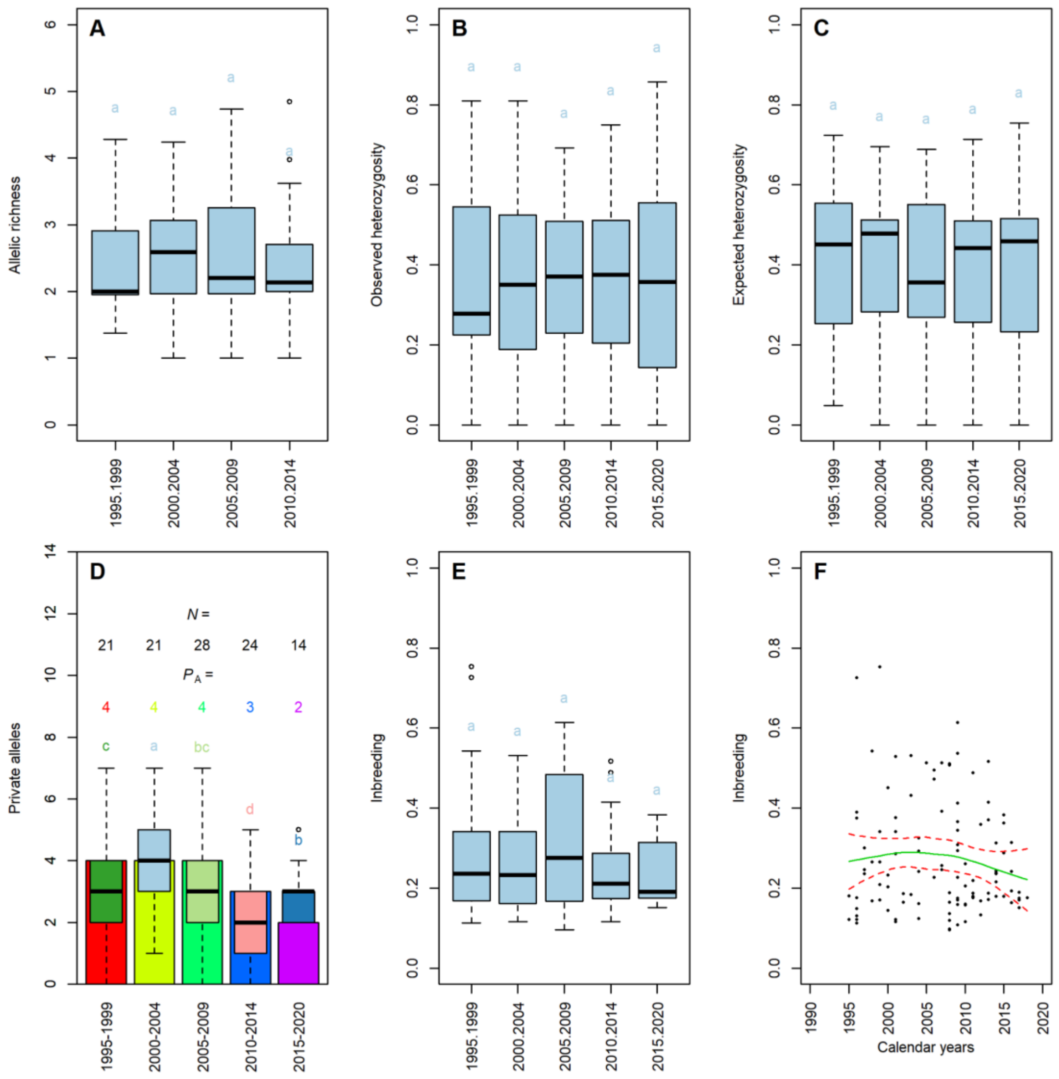

#### Figure S22: Temporal evolution of the east-Med genetic diversity

Boxplot representation of the microsatellites (**A**) allelic richness ($A_R$), (**B**) observed ($H_O$), and (**C**) expected ($H_E$) heterozygosity, (**D**) barplot of the number of private alleles ($P_A$), and boxplot (**E**) and Smooth Local Polynomial Regression Fitting (**F**) representations of the individual inbreeding (*F*) estimates over time. In **A-E** the data is arbitrarily splitted in 5 years time intervals. The letters and boxplot-colors in **A-C** and **E** illustrate the Tukey post-hoc group assignment per period. In **F** the green continuous and red dotted lines represent the Smooth Local Polynomial Regression Fitting and its 95% confidence-interval, respectively.

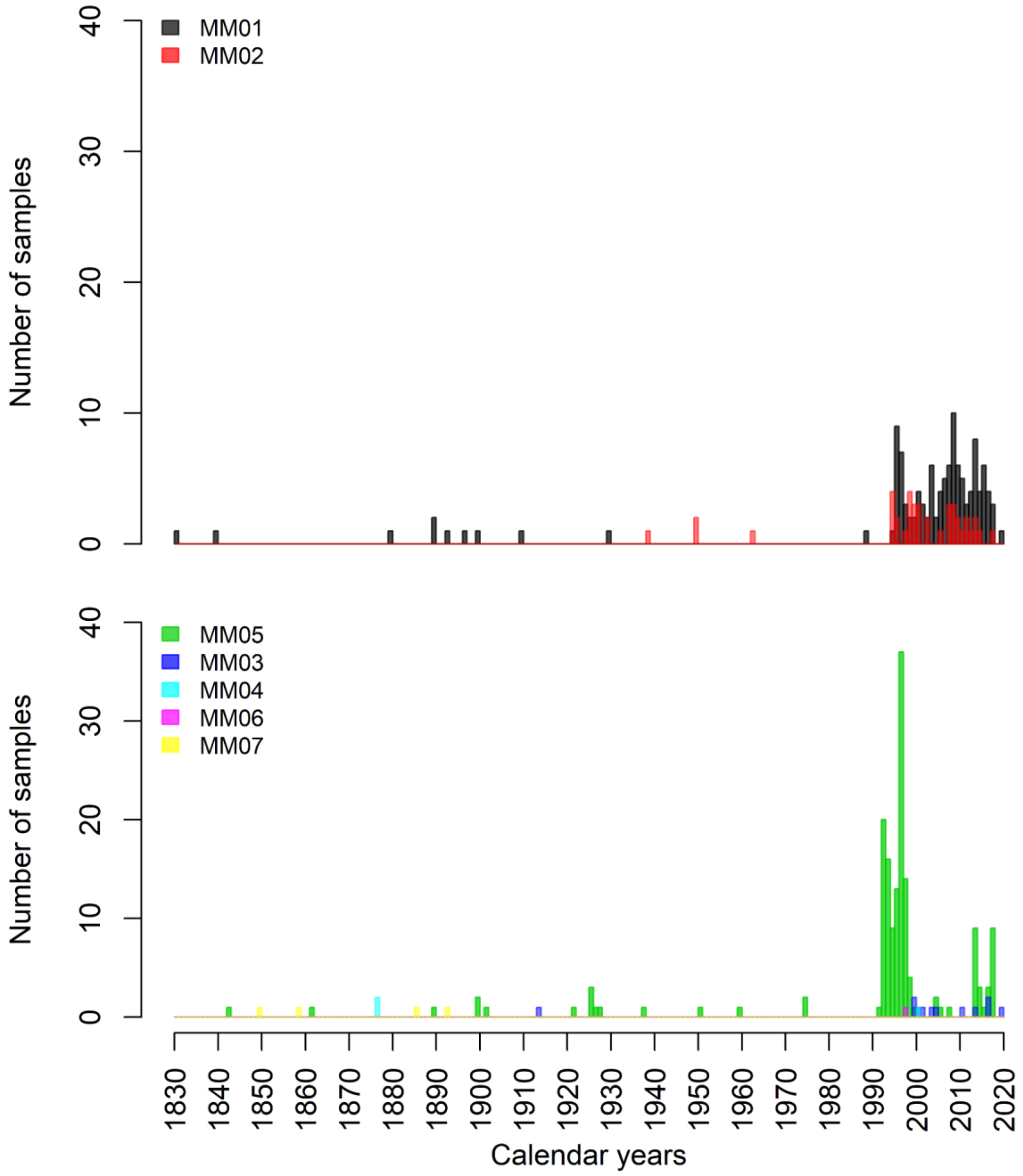


#### Figure S23: MMS d-loop haplotypes distribution over time

The histograms represent the distribution of the d-loop haplotypes included in the present study. For clarity, the haplotypes MM01 and MM02 are represented (upper plot) separately from the others (lower plot). The figure illustrates that the haplotype MM07 has been last seen around 1850 and is very likely to have gone extinct since then, and that the very rare haplotype MM06, last seen in 1998 may have gone extinct by now.

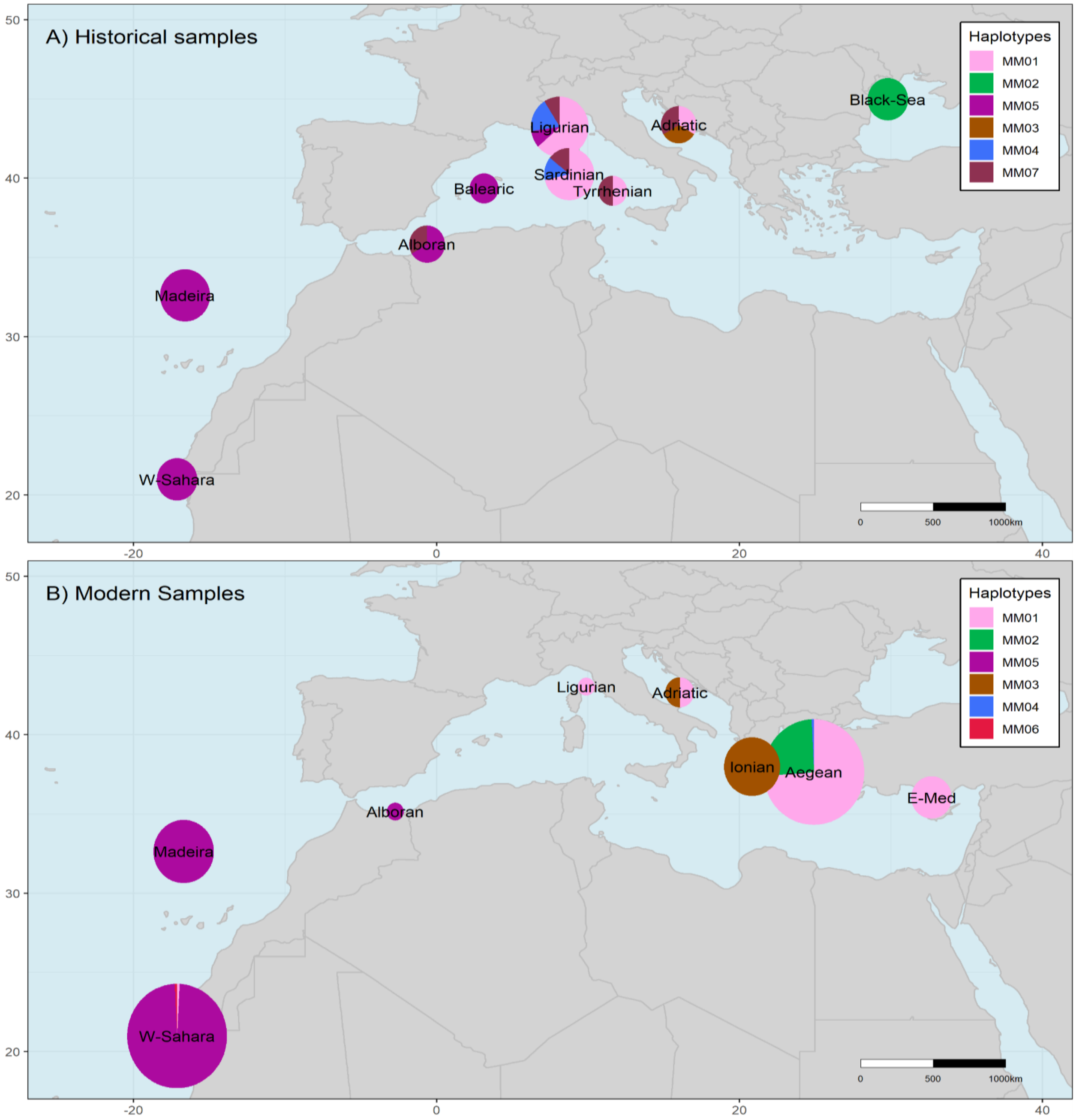


#### Figure S24: MMS d-loop haplotypes geographic repartition over time

The figure presents *M. monachus*' D-loop genetic structure. The historical (**A**) and modern (**B**) haplotype geographic representations illustrate the Mediterranean monk seals temporal changes in mitochondrial genetic diversity in space. The population pie charts show their relative composition in d-loop haplotypes. In both plots, the haplotype names follow (Karamanlidis *et al.*, 2016; Gaubert *et al.*, 2019), and their size is proportional to the log of the number of individuals included (time $5.6e^{-1}$).

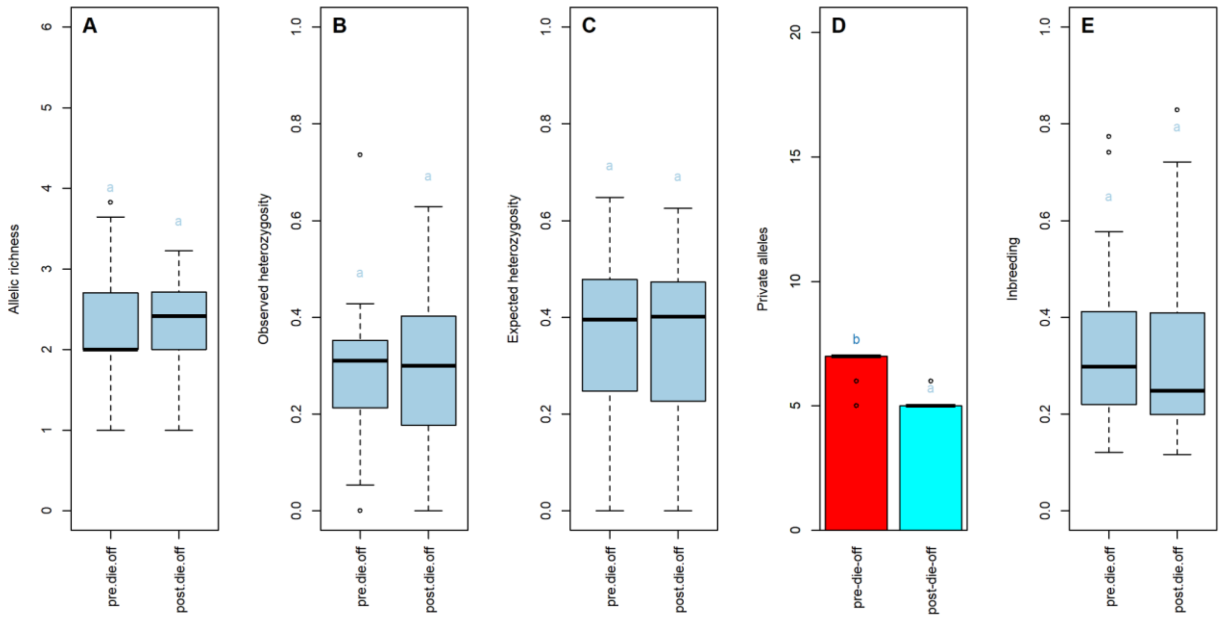


#### Figure S25: Western-Sahara/Mauritania 1997 die-off effect on genetic diversity

Boxplot representation of the microsatellites (**A**) allelic richness ($A_R$), (**B**) observed ($H_O$), and (**C**) expected ($H_E$) heterozygosity, (**D**) barplot of the number of private alleles ($P_A$), and (**E**) boxplot of the individual inbreeding ($F$) estimates, in the Western-Saharan/Mauritania population, before and after the 1997 massive die-off. The letters and boxplot-colors in A-C and E illustrate the Tukey post-hoc group assignment per period. pre/post.die.off: MMS individuals collected and born before / after the 1997 massive die-off event.

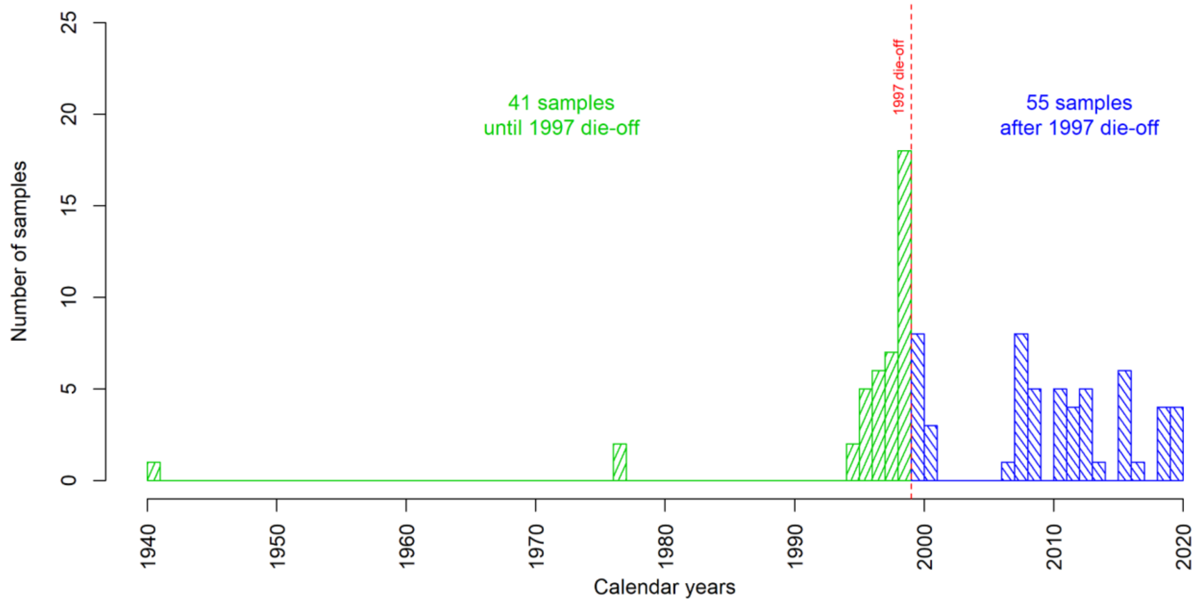


#### Figure S26: Western-Sahara/Mauritania sampling temporality

The figure shows the temporal distribution of genotyped samples included in the present study for microsatellites (**A**) and mtDNA d-loop (**B**) analyses. The lines represent the estimated densities of the respective sampling.

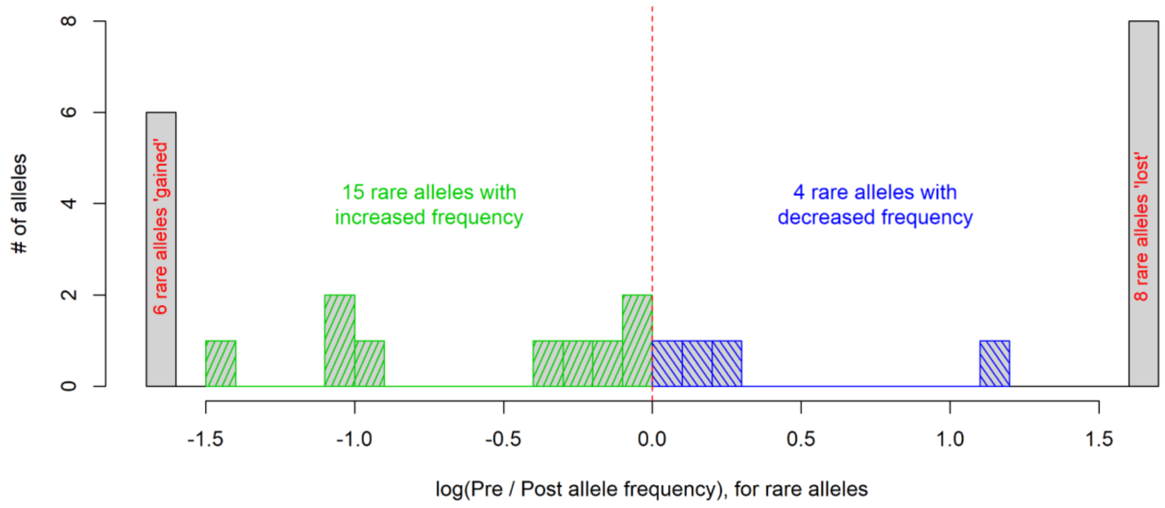


#### Figure S27: Cabo Blanco 1997 die-off effect on rare allele frequencies

Histogram representation of rare allele frequency changes following the 1997 massive die-off. The logarithm of the pre-post ratio of frequencies allows a symmetrical, convenient representation of the increase and decrease in rare allele frequencies. With 15 rare alleles increasing and 4 decreasing in frequency, the 1997 die-off does not appear to have had the expected effect (loss of rare alleles) on the Western-Sahara/Mauritanian population genetic diversity.

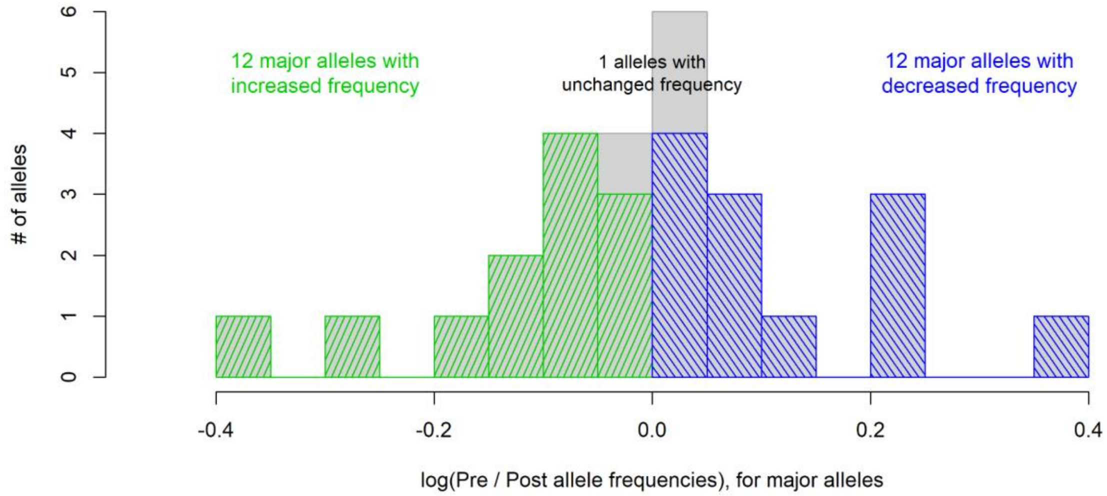


#### Figure S28: Cabo Blanco 1997 die-off effect on major allele frequencies

Histogram representation of major allele frequency changes following the 1997 massive die-off. The logarithm of the pre-post ratio of frequencies allows a symmetrical, convenient representation of the increase and decrease in major allele frequencies. With 12 major alleles increasing and 12 decreasing in frequency, the 1997 die-off does not appear to have had the expected effect (increase of the frequency of major alleles) on the Western-Sahara/Mauritanian population genetic diversity.

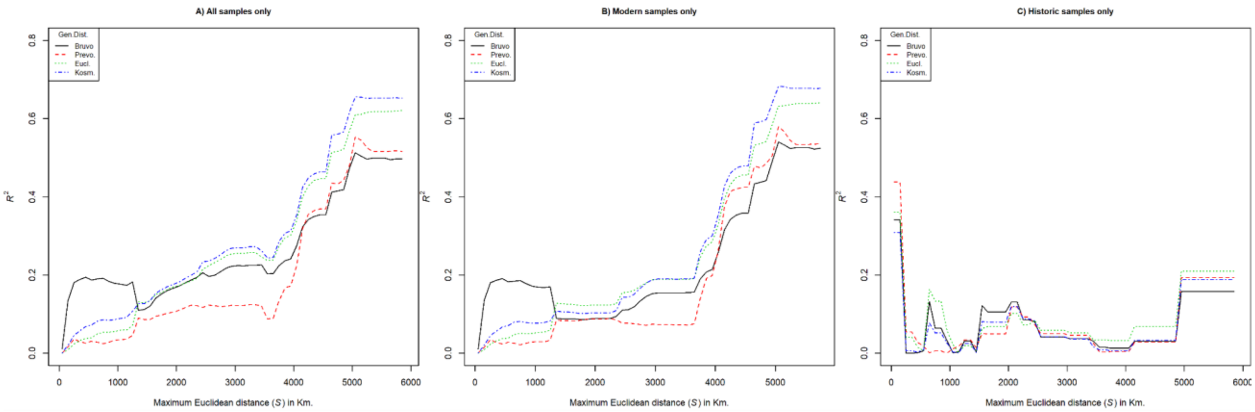


#### Figure S29: Geographic scale influence on IBD

Plot of the variance explained by IBD ($R^2$) against the maximum pairwise geographic Euclidean distance (*S*) estimated from nuclear microsatellite data. Mantel tests are conducted on subsets of pairwise data defined by their maximum geographic distance (*S*) between samples by 100-km increments. The analysis was conducted on all samples (**A**), modern samples (**B**) and historic samples (**C**). For most genetic distances, the best fit to the IBD model is obtained using all the data, i.e. at the largest geographic scale. The highest fit ($R^2$ = 0.7) is found for modern samples only, with Kosman's genetic distances (Kosm) and at a maximum geographic distance (*S*) of 4450 km which corresponds to the entire sampled range of the species. The small sampling size (*N* = 13) and the wide time frame distribution (1800 - 1975) of historic samples alone may partly explain why they do not have a strong IBD signal and why this signal is not much congruent with that of "modern" or "all" sample analyses.

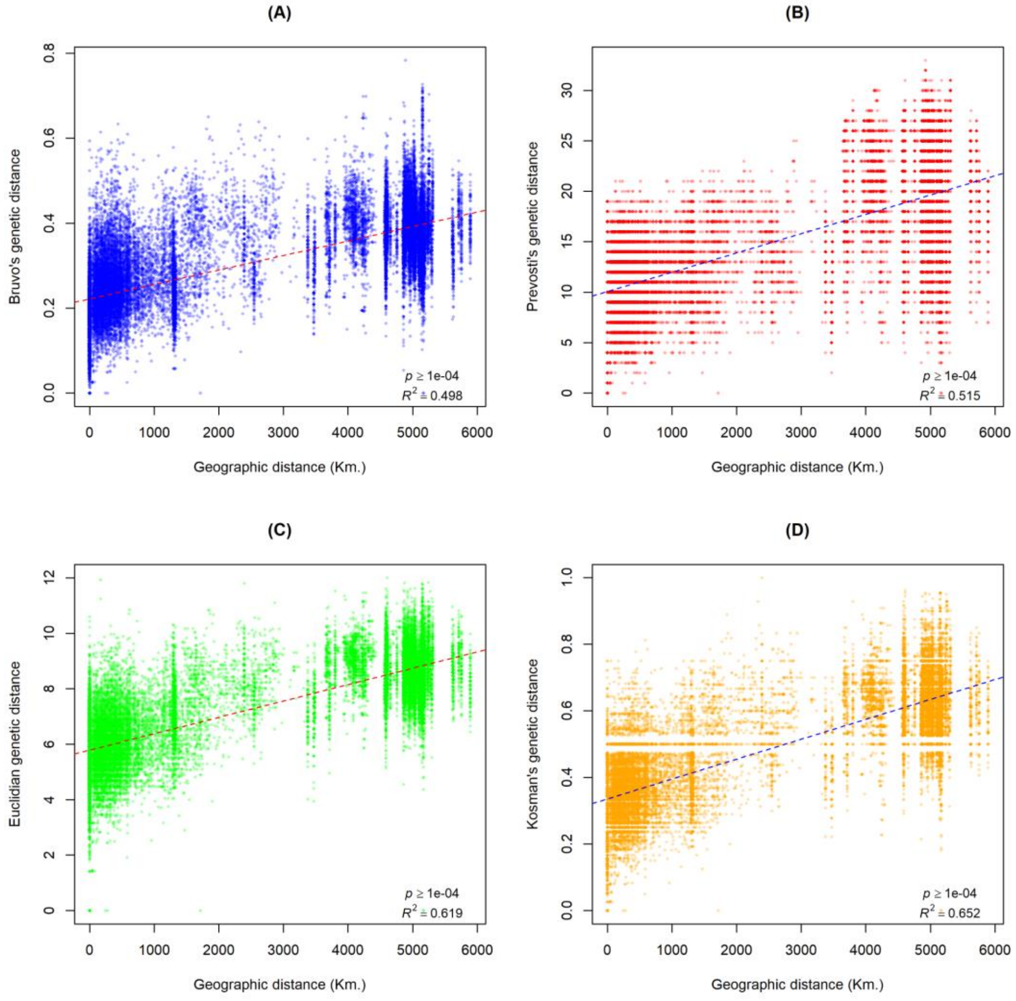


#### Figure S30: Isolation by distance in MMS

Graphic representation of the relationship between geographic and genetic distances (isolation by distance). The individual-based genetic distances estimated from nuclear microsatellite data using Bruvo's **(A),** Prevosti's **(B)**, Euclidian **(C)**, and Kosman's **(D)** genetic distances at their highest relationship with geographic distance (Fig. S_IBD.S.all) are represented against geographic distance. The linear model (dashed lines), the Mantel tests explained variance ($R^2$) and *p*-value (*p*) are shown to support the represented trends. The figure shows a positive relationship of nuclear data with Euclidean geographic distances suggesting that the gene flow may be impacted by geographic features.

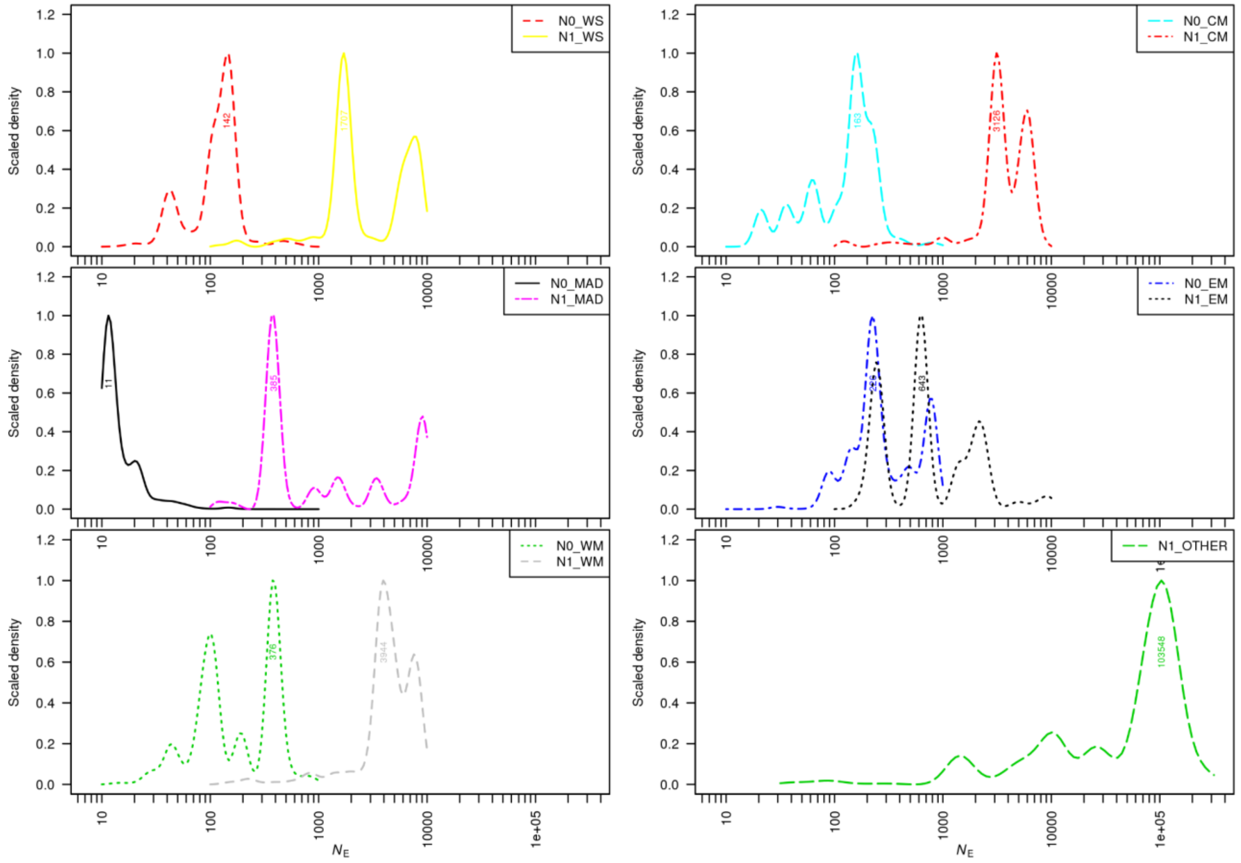


#### Figure S31: ABC population size posteriors

ABC-GLM posterior scaled-density distributions of the effective population size ($N_e$) under the model 181, represented after ($N_0$: current population) and before ($N_1$: ancient population) the demographic event occurring at $T_1$ (Fig. X). The posterior distributions suggest that all populations underwent major declines, most of which of at least one order of magnitude. Furthermore, the several estimates of the current population ($N_0$) ecompasses highly realistic values including those of recent census data (e.g. Madeira & Western-Sahara/Mauritania). WS: Western-Sahara/Mauritania, MAD: Madeira, WM: Western-Mediterranean, CM: Central Mediterranean, EM: Eastern-Mediterranean.

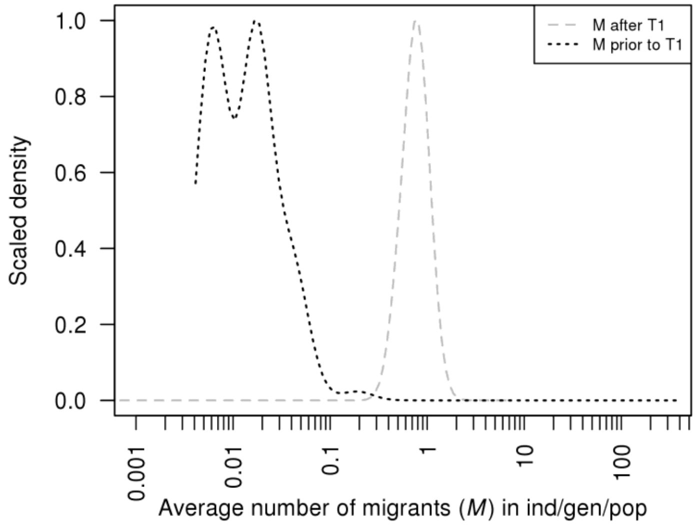


#### Figure S32: ABC migration rate posteriors

ABC-GLM posterior scaled-density distributions of the migration rate ($M_i$) under the model 181, represented after ($M_0$: current population) and before ($M_1$: ancient population) the demographic event occurring at $T_1$ (Fig. X). The posterior distributions suggest that after the event occurring at $T_1$ (that includes population size decrease, loss of population and change in connectivity), the sampled population needs a high migration rate to fit the model. Contrastingly, before the loss of connecting populations, low migration rates were sufficient to ensure connectivity among colonies/populations.

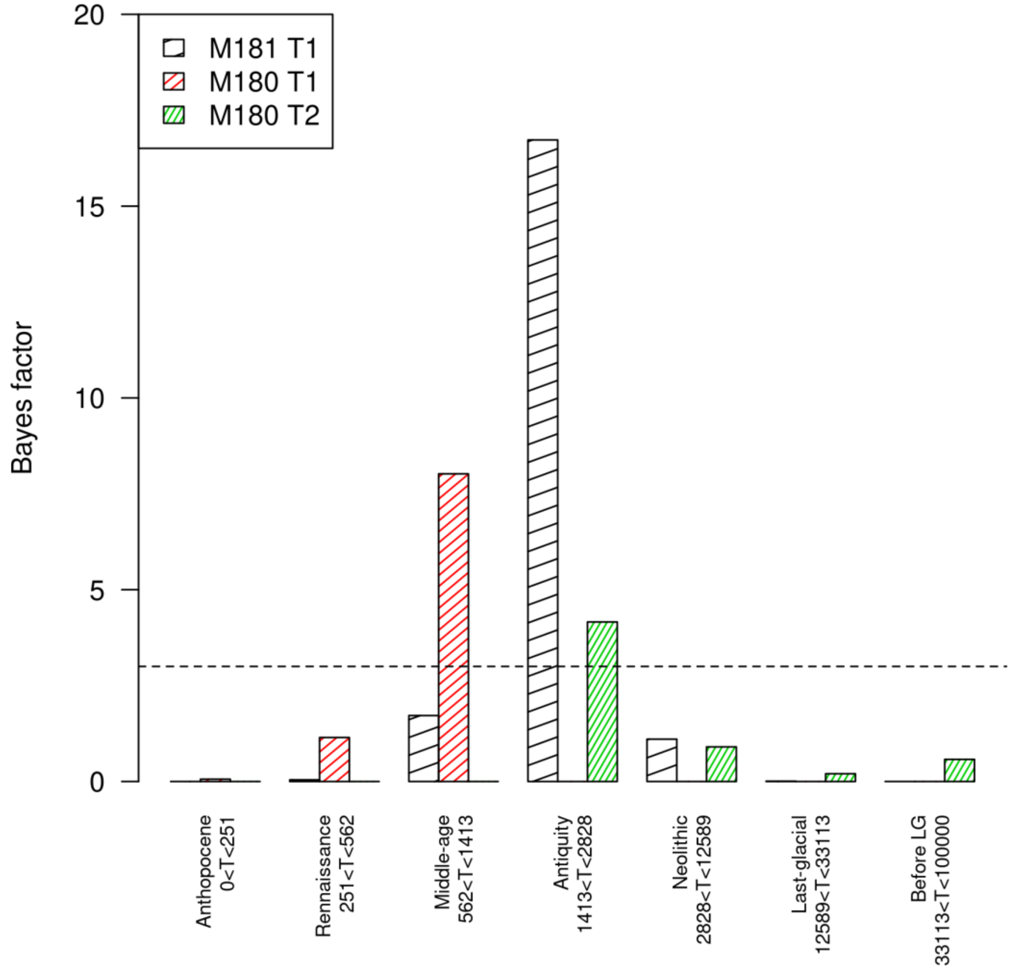


#### Figure S33: Bayes factor of period of MMS' decline.

Most likely periods of the species decline. Bayes facor values for seven alternative time period for the decline of the species' populations and the fragmentation of their habitat, estimated from the two most supported models.
